# Monkeypox virus shows potential to infect a diverse range of native animal species across Europe, indicating high risk of becoming endemic in the region

**DOI:** 10.1101/2022.08.13.503846

**Authors:** Marcus SC Blagrove, Jack Pilgrim, Aurelia Kotsiri, Melody Hui, Matthew Baylis, Maya Wardeh

## Abstract

**Background:** Monkeypox is a zoonotic virus which persists in animal reservoirs and periodically spills over into humans, causing outbreaks. During the current 2022 outbreak, monkeypox virus has persisted via human-human transmission, across all major continents and for longer than any previous record. This unprecedented spread creates the potential for the virus to ‘spillback’ into local susceptible animal populations. Persistent transmission amongst such animals raises the prospect of monkeypox virus becoming enzootic in new regions. However, the full and specific range of potential animal hosts and reservoirs of monkeypox remains unknown, especially in newly at-risk non-endemic areas.

**Methods:** Here, utilising ensembles of classifiers comprising different class balancing techniques and incorporating instance weights, we identify which animal species are potentially susceptible to monkeypox virus. Subsequently, we generate spatial distribution maps to highlight high-risk geographic areas at high resolution.

**Findings:** We show that the number of potentially susceptible species is currently underestimated by 2.4 to 4.3-fold, and that a high density of wild susceptible species are native to Europe. We provide lists of these species, and highlight high-risk hosts for spillback and potential long-term reservoirs, which may enable monkeypox virus to become endemic.

**Interpretation:** We highlight the European red fox and brown rat, as they have established interactions with potentially contaminated urban waste and sewage, which provides a mechanism for potential spillback. We anticipate that our results will enable targeted active surveillance of potential spillback event, to minimise risk of the virus becoming endemic in these regions. Our results also indicate the potential of domesticated cats and dogs (latter now confirmed) being susceptible to monkeypox virus, and hence support many health organisations’ advice for infected humans to avoid physical interaction with pets.

## Introduction

The largest ever outbreak of monkeypox virus (MPXV), outside of endemic countries, was first identified in May 2022^1^. Previously a rare zoonotic virus, this ongoing outbreak has so far resulted in over 35,000 confirmed cases across all six major continents, causing the World Health Organisation to declare it the 7^th^ ‘Public Health Emergency of International Concern’ in July 2022. Whilst only a small number of deaths have occurred so far outside of endemic regions, the incidence is still steadily increasing despite initial mitigation strategies including vaccination, contact tracing^2^ and voluntary isolation^3^.

As is common with emerging zoonotic viruses, the animal host range is central to the epidemiology of the virus. The host range affects the risk of spillover into human populations, dissemination of virus to new regions (by movement of infected hosts), and the establishment of novel reservoirs of the virus – sustaining it between outbreaks and making eradication more difficult or impossible^4^.

The natural reservoirs of monkeypox virus are likely a range of African rodents, including tree squirrels, rope squirrels, Gambian poached rats, as well as various species of monkeys^5^. In the 2003 Midwest USA outbreak, MPXV was imported via three genera of African rodents (*Cricetomys, Graphiurus, Funisciurus*)^6^. Once in the USA, these rodents were housed with and subsequently infected prairie dogs (*Cynomys ludovicianus*), with whom direct contact produced all human cases in this outbreak.

Outbreaks, such as 2003, have demonstrated that animal species not found in the MPXV-endemic region can serve as hosts and transmit the virus to humans. However, the full extent of the host range of MPXV remains unknown. This is of particular concern in non-endemic regions, where local hosts which have never been studied for MPXV competence could serve as reservoirs following spillback from humans into animal populations. Should animal species outside of the current endemic region be susceptible to the virus, and spillback into these susceptible species occur, the current MPXV outbreak could be prolonged or even become endemic in new regions. This concern is paramount in the current (2022) outbreak of MPXV, where the virus has persisted in non-endemic regions for considerably longer and is far more geographically widespread than in any previous record. Furthermore, the first recorded instance of human-to-animal transmission occurred very recently, with the infection of a pet dog in France^7^ – this demonstrates that there is a real and immediate risk of human-to-animal transmission, and that the need to understand the full potential host range is critical to ongoing mitigation efforts.

In addition to MPXV, there are many poxviruses of major human and animal health concern. These include the now eradicated smallpox, the zoonotic cowpox^8^ and orf^9^ viruses, the ecologically pivotal squirrelpox and myxoma viruses, and the economically important pseudocowpox and bovine papular stomatitis viruses^10^.

Similarly to monkeypox, the known host ranges of other poxviruses are understudied and very varied. For example, many poxviruses have only a single known host species, whilst the well-studied cowpox virus has 70 known (observed) host species. This large knowledge gap, for both MPXV and other poxviruses, needs to be resolved in order to understand the spillover, spillback and reservoir risks of these highly important human and animal pathogens.

To this end, we deploy ensembles of lightGBM (gradient boosting decision trees) models, incorporating instance weights (to proxy research effort), and a range of class balancing techniques (to correct for class imbalance), to predict which animal species are potential hosts of each poxvirus. Our approach integrates data from three perspectives encompassing: (1) genomic features depicting various aspects of poxviruses extracted from all available complete genomes (63 species); (2) phylogenetic, taxonomic, ecological, environmental and geospatial traits of mammalian (n⍰=⍰1,489) and avian species (n = 995); and (3) topological characteristics describing the linkage of poxviruses with their observed and potential hosts, and of current knowledge of all viral sharing amongst mammals and birds (24,445 interactions).

Using this methodology, we are the first to provide detailed predictions of susceptible hosts specifically for poxviruses. We show that the number of potential host species for monkeypox virus is currently underestimated by 2.4 to 4.3-fold, by 2.34 to 4.56-fold for mammalian poxviruses, and by 1.96 to 4.02-fold for avian poxviruses.

Of particular note, our results show that there are many potentially MPXV-susceptible host species across Europe, and of particular concern for the virus potentially becoming endemic in this region are the red fox (*Vulpes vulpes*) and the brown rat *(Rattus norvegicus*). The red fox frequently scavenges from domestic waste and could become infected via contaminated fomites, and brown rats inhabit sewerage systems across Europe and could become infected by faecal shedding. The data, predictions and geographic ranges presented here will enable more efficient triage of potential hosts for surveillance programmes, and hence focused risk estimation, and mitigation procedures.

## Results

### Summary of predictions for monkeypox virus

Our pipeline to predict associations between poxviruses and their hosts indicated a total of 222 mammalian species in which MPXV has been observed or could be found at mean probability threshold⍰of >0.5, with standard deviation (SD) of −48/+59; and 129 at mean probabilitythreshold⍰of ≥0.7602445 with SD of −18/+46. For simplicity, hereafter we refer to these results in the following format: The number of hosts associated with MPXV is >0.5=222 (−48/+59); ≥0.76=129(−18/+46). See methods for explanation of ‘0.76’ threshold.

Hence, in addition to the 51 observed MPXV host species, the number of new predicted mammalian species in which MPXV has not been observed to date is >0.5=171 (−48/+59); ≥0.76=78(−18/+46). Figure 1 illustrates the predicted wild species broken down by taxonomic order (Figure 1-A), as well as top 20 predictor groups with most influence (mean SHAP value) on predicted association between mammals and MPXV (Figure 1-B), and top 10 predicted new host species (ranked by mean probability) in each of the following orders: Primates, Rodentia and Carnivora (Figure 1-C). Supplementary Data 1 lists full results for MPXV.

**Figure 1.**
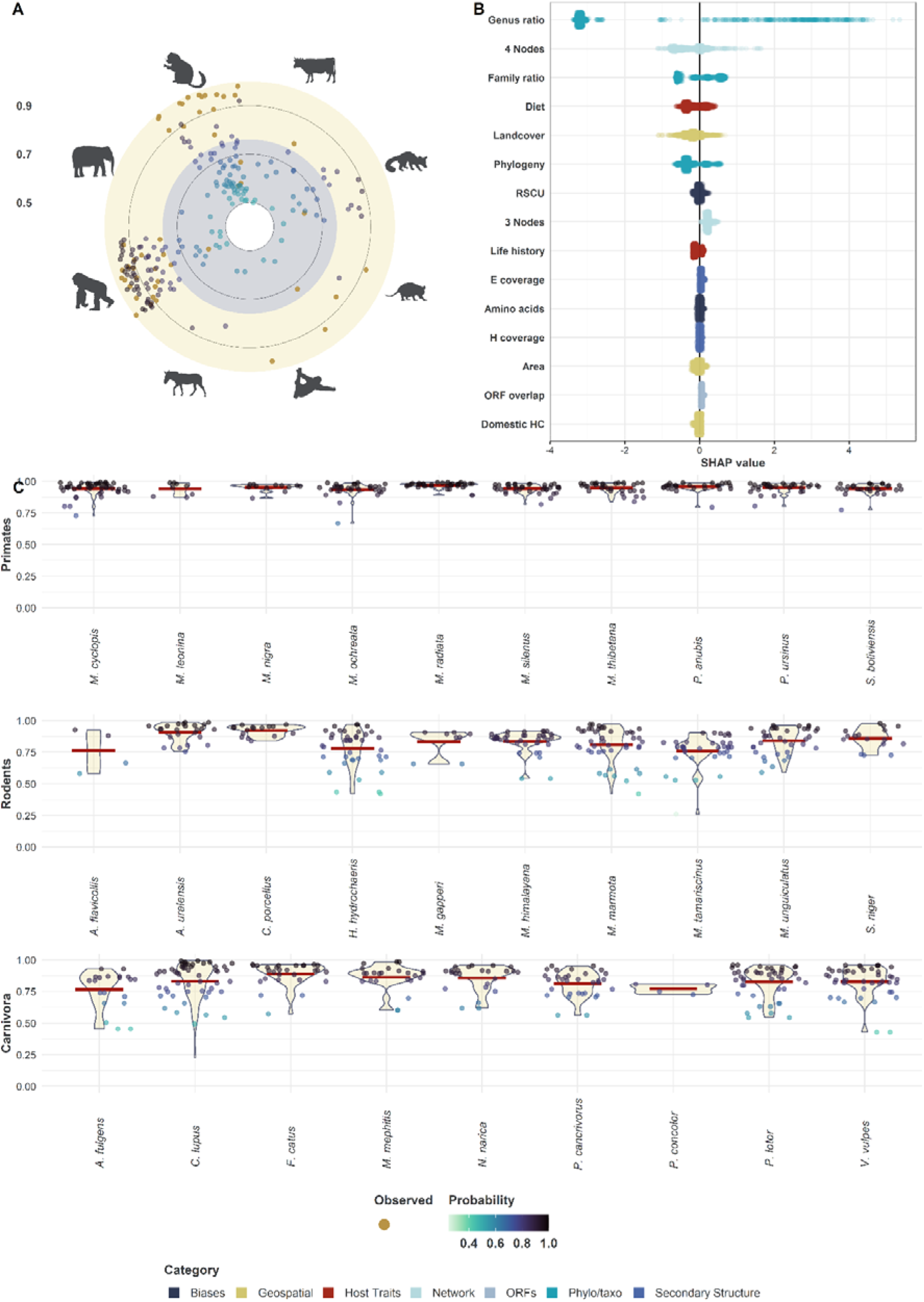
Monkeypox Virus (MPXV) Results. **Panel A: MPXV hosts**. Yellow points represent observed MPXV hosts. Light green to dark blue points represent new host species predicted by our top 25 ensembles (each comprise 4 constituent models trained with the following class balancing techniques: SMOTE, SMOTE (25%), SMOTE-ENN, and SMOTE-ENN (25%), see methods section for explanation of balancing techniques). The grey area corresponds to mean probability >0.5 and <0.76, yellow area corresponds to mean probability ≥0.76. The Y-axis represents the mean probability produced by highest performing 25 ensembles (100 models in total) – ranging from 0 to 1. The X-axis represents the order of the host, as follows (clockwise): Artiodactyla, Carnivora, Didelphimorphia, Other (mammals), Perissodactyla, Primates, Poboscidea, and Rodentia. The percent of observed MPXV (known) associations predicted by the final model (mean probability of the top 25 ensembles) were as follows: >0.5=0.98(−0.039/0); ≥0.76=0.863(−0.118/+0). **Panel B: SHAP plot of the top 20 feature groups**. SHAP values were calculated for all possible associations of MPXV and all mammalian species included in our models (n=1,486), for each constituent model (n=4) in our ensembles (n=25). SHAP values were averaged for each association/feature combination, and then summed for each association/group combination. Each point corresponds to MPXV-mammalian associations and is colour-coded by standardised feature value. Features are colour coded by their category. **Panel C: Top predicted MPXV host species in three orders – Primates (n=10), Rodentia (n=10), Carnivora (n=10)**. Newly predicted (previously unobserved) species were ordered by mean probability of the top 25 ensembles, and top 10 species were selected from each order. Points represent predictions (probability) of each constituent model in each ensemble and are coloured by probability. Violin plots show the kernel probability density of predictions (probability) at different values, and horizontal red lines represent overall mean probability (as utilised in selection).

### MPXV Predictors

Taking our top 25 ensembles (4 models each), the top 10 predictor groups (by summation of absolute mean SHAP values) of whether a given mammalian species could be susceptible to MPXV are as follows: Genus ratio =3.993 ±0.669, 4 nodes (network) = 1.875±0.292, family ratio = 1.497±0.189, diet = 1.456±0.133; landcover =1.433±0.182; phylogeny = 1.357±0.123; codon bias = 1.326±0.049; 3 nodes (network) = 1.255±0.054; life history = 1.139±0.044; and E coverage (secondary structure) = 1.112±0.032. For precise definitions and equations of predictors see methods and supplementary materials.

### Summary of predictions for all poxviruses

Overall, our pipeline predicts >0.5=1,251 (−379/+ 415); ≥0.76=576(−169/+370) previously unobserved associations that could potentially exist between >0.5=422(−56/+40); ≥0.76=277 (−60/+90) mammalian species and 0.5=45(−6/+3); ≥0.76=35(−5/+4) poxviruses (species or strain). For avian species, our results are as follows: >0.5= 156 (−47/+50); ≥0.76= 91(−27/+37) previously unobserved associations between >0.5=89(−17/+4); ≥0.76=64(−14/+13) avian species and 0.5=11 (−2/+0);≥0.76=6(−2/+4) poxviruses. Figure 2 visualises observed and predicted associations. Supplementary Data 2 and 3 list full results for mammalian and avian poxviruses, respectively.

**Figure 2.**
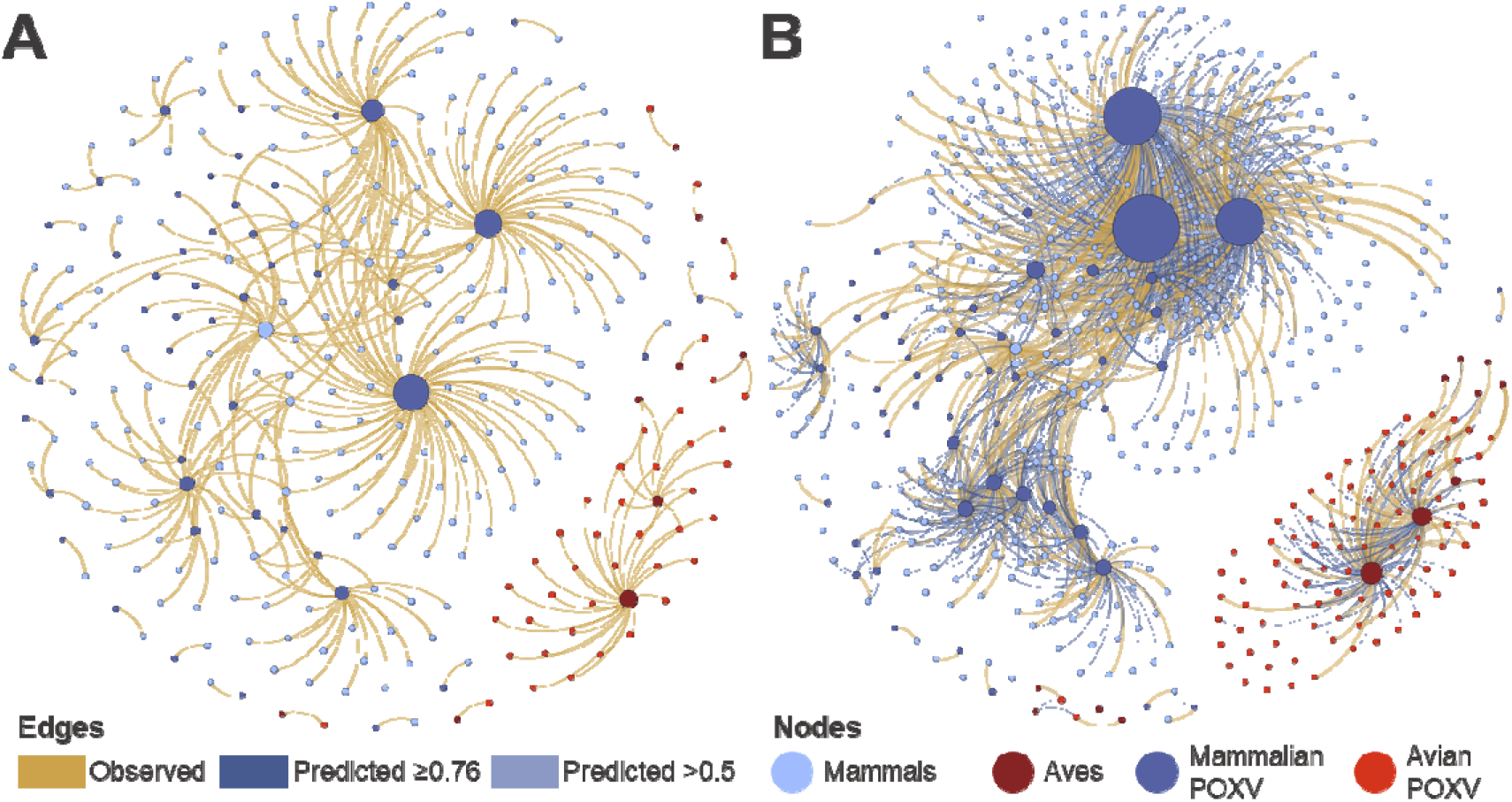
Observed and predicted poxvirus-susceptible species network. **Panel A – Potential Motifs in virus-host networks**. **Panel A – Network of observed poxvirus-mammal/aves associations**. Edges represent observed poxvirus-mammal/aves associations (n=362). Nodes represent poxviruses with at least one observed avian/mammalian host species (n=60), and their observed mammalian (n= 216) or avian (n=41) hosts. Nodes are coloured by category (mammal, aves, mammalian poxvirus, avian poxvirus), and are sized in proportion to number of poxviruses/susceptible species observed per node. **Panel B – Network of observed and predicted poxvirus-mammal/aves associations**. Nodes represent poxviruses included in this study, and their observed or predicted mammalian (n=446) or avian hosts (n=91). Nodes are coloured by category (mammal, aves, mammalian poxvirus, avian poxvirus). Yellow edges represent observed associations (n=362), light blue edges represent associations predicted at probability threshold >0.5 and <0.76 (n=740), and dark blue edges represent associations predicted at threshold ≥0.76 (n=667). Thickness of edges is in proportion to their probability (observed association = 1; predicted association = mean ensemble probability).

On average, each of >0.5= 537(−42 /+27); ≥0.76=442(−37/+53) species in which poxviruses has been observed or predicted by our pipeline, is associated with >0.5= 3.294 (−0.581/+0.667); ≥0.76=2.328(−0.674/+1.939) poxviruses. In wild species (>0.5= 508(−42/+27); ≥0.76=413(−37/+53)) these results were as follows: >0.5=2.947(−0.522/0.638); >=0.76=2.051(0.617/1.803). Figure 3-A illustrates the top predicted wild species, by number of observed and predicted poxviruses, in selected host orders.

**Figure 3.**
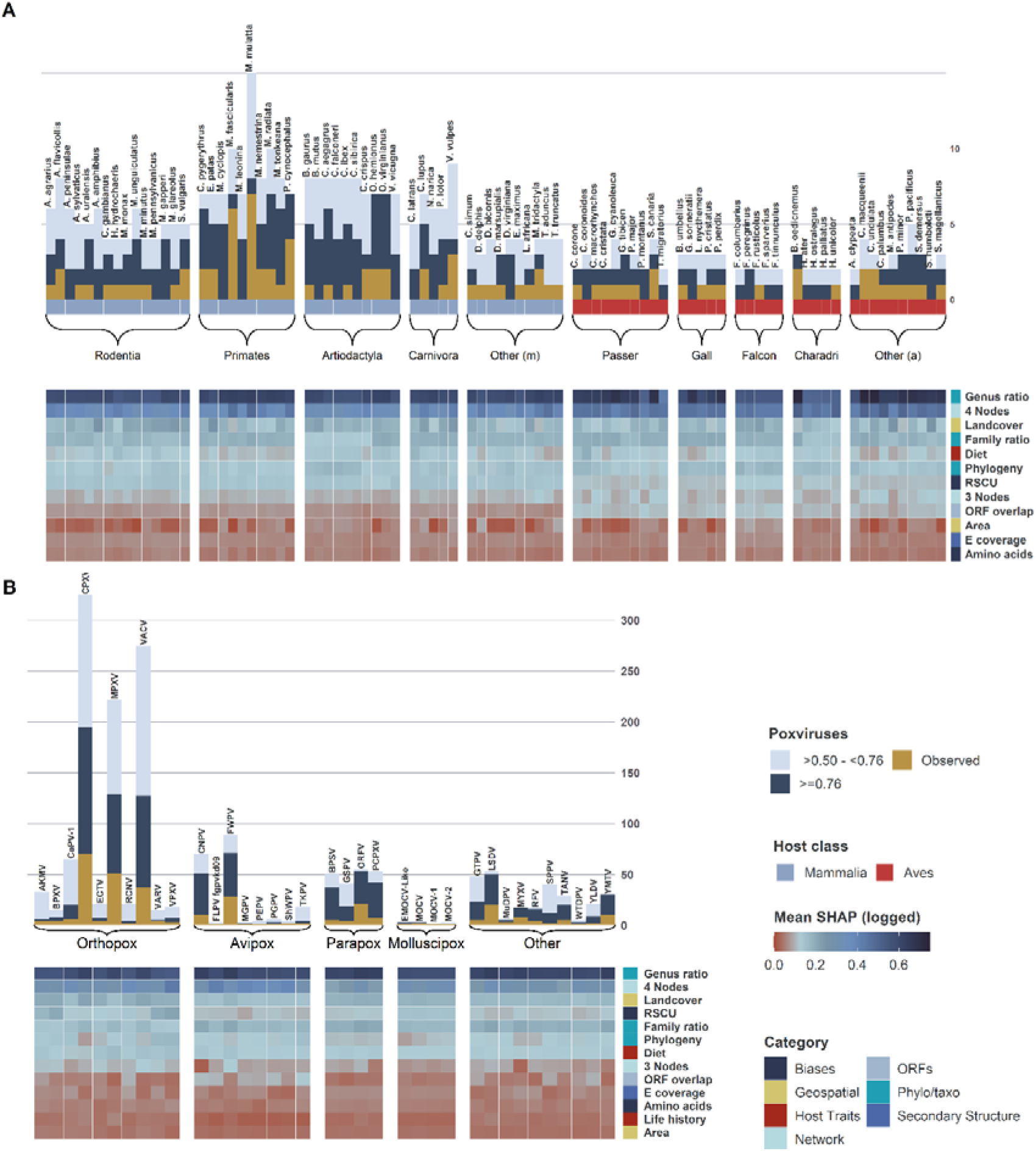
Top susceptible host species and poxviruses. **Panel A: Number of poxviruses per susceptible wild species**. Susceptible wild species were grouped by class (mammalian or avian) and order; orders were arranged by number of species with at least one observed or predicted poxvirus. The top species (by total number of poxviruses = observed + predicted) were selected as follows: Rodentia (n=15), Primates (n=10), Artiodactyla (n=10), Carnivora (n=5), other mammalian orders (n=10); for aves, Passeriformes (n=10), Galliformes (n=5), Charadriiformes (n=5), Falconiformes (n=5), and other avian orders (n=10). Yellow bars represent number of poxviruses observed to be found in each susceptible species. Blue stacked bars show other poxviruses predicted to be found in each species by our pipeline. Predicted poxviruses per host are grouped by association probability into two categories: dark blue = ≥0.76, and light blue = >0.5 - <0.76. Heatmaps represent log_10_ mean absolute SHAP values. SHAP values were computed separately for each possible association of each of the included animal and poxvirus (n=63), for each constituent model (n=4), of our top ensembles (n=25). Mean absolute SHAP values were taken for each association/feature combination, and then summed for each association/group combination. Features are colour coded by their category (see methods). Supplementary Figure 4 visualises results for humans and domesticated animals. **Panel B: Number of susceptible wild species per poxvirus**. Poxviruses were grouped by their genus, and ordered by the number of poxviruses with at least one observed and/or predicted susceptible species. Top species (by total number of susceptible species = observed + predicted) were selected as follows: Orthopoxvirus (n=15), Avipoxvirus (n=8), Parapoxvirus (n=4), Molluscipoxvirus (n=4), and other poxviruses (n=10). Yellow bars show the number of poxviruses observed to be found in each species. Blue stacked bars represent other poxviruses predicted to be found in each host by our pipeline. Predicted poxviruses per host are grouped by association probability into two categories: dark blue = ≥0.76, and light blue = >0.5-<0.76. Heatmaps represent log_10_ mean absolute SHAP values. SHAP values were computed separately for each possible association of each included virus, and all possible mammalian (n=1,489) or avian (n=995) hosts, for each constituent model (n=4), of our top ensembles (n=25). Mean absolute SHAP values were taken for each association/feature combination, and then summed for each association/group combination. Features are colour coded by their category.

Our results indicate that the average host range of the 63 fully sequenced poxviruses (*species or strain*) which entered our model is: >0.5= 28.079 (−6.762/+7.381); ≥0.76= 16.333 (−3.111/+6.460). Figure 3-B illustrates the top predicted poxviruses, by number of observed and predicted hosts, in selected genera.

### Predictors of Mammalian and Avian poxviruses

Genus ratio, 4 nodes (networks) and family ratio were the top three predictor groups for both mammalian (2.826 (±0.462); 0.864 (±0.47); 0.477 (±0.129)) and avian poxviruses (2.968 (±0.393); 0.86 (±0.474); 0.532 (±0.08)), respectively. Phylogeny and landcover was the most important predictor group from the host perspective for mammals (0.291 (±0.171); 0.282 (±0.21)) and birds (0.288 (±0.113); 0.256 (±0.188)). From the virus perspective, codon biases were the most important predictor group for mammalian poxviruses (0.164 (±0.12)) – ranked 8^th^ in total), whereas ORF overlap was the most important for avian poxviruses (0.176 (±0.083) – 6^th^ in total).

### Mapping predictions

Spatial distribution of observed host species as well as those predicted as susceptible were visualised by summarising IUCN^11^ and Birdlife^12^ range maps for terrestrial mammals and birds into a grid with cells measuring 1/6 × 1/6 of a degree. Figure 4 visualises global distribution of MPXV (observed=47/ predicted & observed = 190 species), wild mammalian (176/384 species), and wild avian (29/71 species) species susceptible to poxviruses (41 and 12 viruses, respectively). See Supplementary Figures 5-7 for host order-level global distributions.

**Figure 4.**
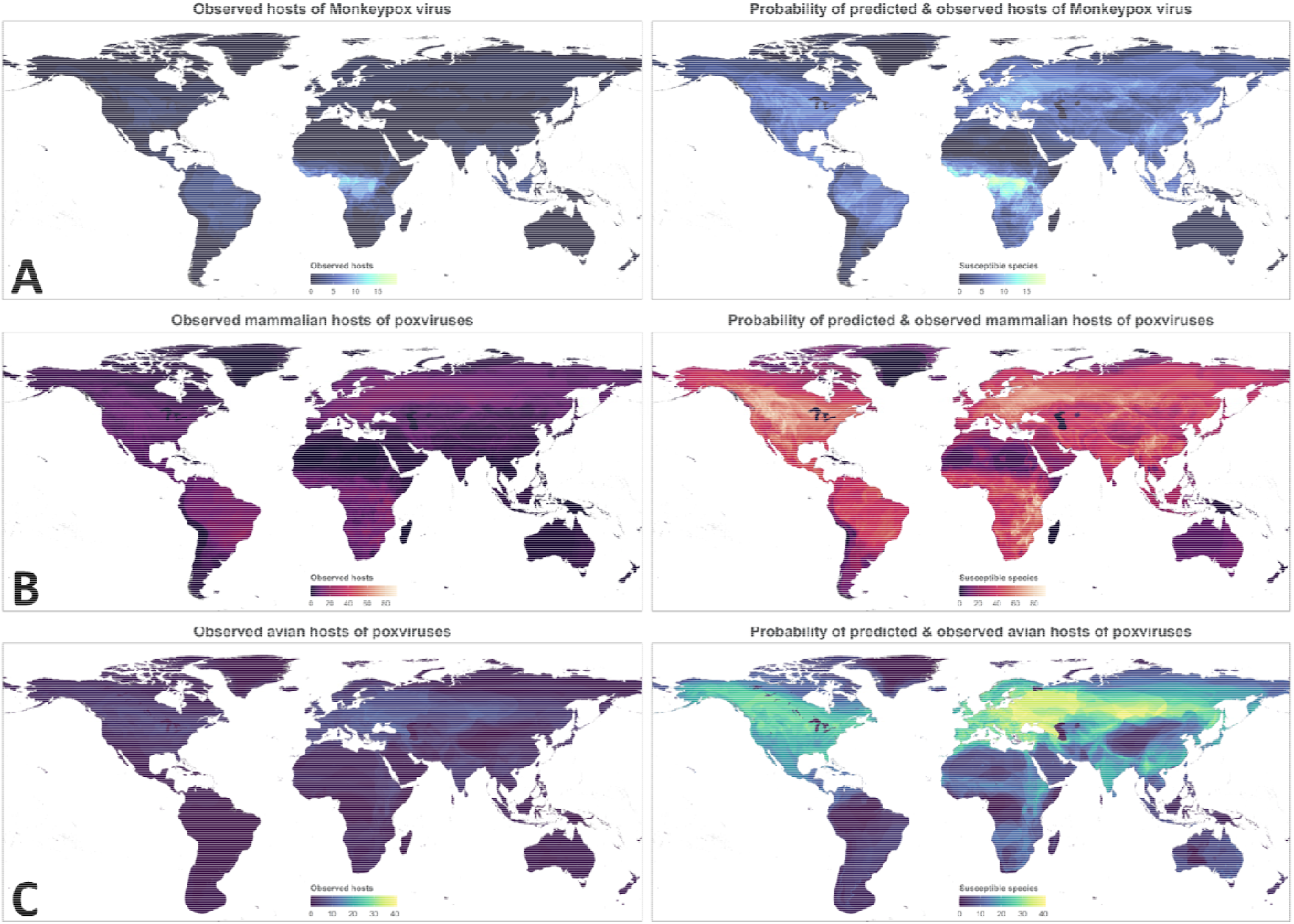
Global distribution of observed and predicted susceptible wild terrestrial species. **Panel A – Monkeypox virus. Panel B – Mammalian poxviruses. Panel C – Avian poxviruses**. Rasters were computed from presence shapefiles of observed (known) species (left-hand side) by summing number of species per each grid cell (each measuring cells measuring 1/6 × 1/6 of a degree). The right-hand side shows both observed host species and predicted susceptible species (threshold >0.5). Each grid cell is coloured by the sum of probabilities of all predicted hosts (observed host = 1; predicted host = square root of prediction value). The colour scales for both maps in each row is the same to allow comparison.

### Model performance

We evaluated the performance of our pipeline against held-out test sets (stratified random samples comprising 10% of all possible associations, n= 87,532). Over 100 iterations, our ensemble of four class balancing techniques (SMOTE, SMOTE (25%), SMOTE-ENN, and SMOTE-ENN (25%)) achieved the following: AUC = 0.994 (SD=+/−0.002); PR-AUC =0.485 (+/−0.081); F1-Score – >0.5=0.309 (+/−0.04); ≥0.76=0.422 (+/−0.056); TSS – >0.5=0.909 (+/−0.049); ≥0.76=0.827 (+/−0.073)); NPV – >0.5=0.999 (+/−.0002); ≥0.76=0.999 (+/−.0003); PPV (precision) – >0.5=0.186 (+/−0.029); ≥0.76=0.285 (+/−0.051); sensitivity (recall) – >0.5=0.926 (+/−0.05); ≥0.76 = 0.836 (+/−0.074); and specificity – >0.5=0.982 (+/−0.004); ≥0.76 = 0.991 (+/−0.003). Supplementary Figures 8-11 illustrate performance metrics calculated per each constituent model for all ensembles (n =100) and top performing ensembles (n=25).

## Discussion

Currently, as of August 2022, we are experiencing the largest and longest lasting outbreak of the zoonotic monkeypox virus. The persistence of the virus outside of its endemic region raises new threats of it becoming endemic via increased opportunity to spillback into wild and domesticated animal populations. Until now, the range of potentially susceptible host species has not been defined. Here, using ensembles of classifiers comprising different class balancing techniques and incorporating instance weights, we provide the first specific and comprehensive poxvirus–animal host susceptibility predictions. We identify multiple-fold underestimations in current understanding of potential monkeypox virus – and other poxvirus – hosts. For monkeypox specifically, we show high probability of susceptibility to MPXV amongst a number of wild species across Europe, which aligns with the current outbreak epicentre, and hence highlight the immediate risk of spillback and potentially endemicity. This work will enable more efficient triage of potential hosts for surveillance programmes, and hence and focused risk estimation, and mitigation procedures.

### Monkeypox Virus

Our study highlights a current underestimation of the animal host range of monkeypox virus of between 2.4- and 4.3-fold. Approximately 80% of the newly predicted hosts were from the Rodentia and Primates orders. Our improved estimation of the potential host range of this virus enables us to better understand, predict and mitigate potential spillback events into animals and thus enable policy makers and surveillance strategies to minimize the risk of monkeypox virus becoming endemic, via animal reservoirs, in new regions.

The high density of predicted monkeypox virus hosts in south-east European region of Hungary/Romania, and to a lesser extent Ukraine/Belarus, raises the concern of potential spillback into susceptible wild hosts.

Of particular note, the brown rat (*Rattus norvegicus*), a known host of cowpox virus and extremely common throughout European sewerage systems is also a predicted MPXV host. As a high proportion of patients shed MPXV through their faeces^13^, and MPXV DNA has been detected in wastewater^14^, there is a clear route to infection of brown rat populations. In addition, the urbanized red fox (*Vulpes vulpes*) was also a high probability susceptible host, and, with this species’ close association to both humans and rodents, represents a high-risk bridge-species to facilitate inter-species viral sharing. The red fox is an urban scavenger, hence, is likely to come into contact with contaminated household waste fomites – providing a route to infection of the species. Our study highlighted other European rodents, including the herb field mouse *(Apodemus uralensis*), Alpine marmot (*Marmota marmota*) and the yellow-necked field mouse (*Apodemus flavicollis)*. All three of these species are native to Europe – in particular the high-risk region defined here – and have pockets of very high population density, making them ideal long-term reservoir hosts. Given the current disproportionately high number of human cases across Europe^1^, our findings suggest that these species should be surveillance priorities.

The phylo/taxo category were the top predictors for MPXV susceptibility of the above mice, marmot, fox, and brown rat, indicating that predicted susceptibility to MPXV in these key species is strongly linked to their phylogenetic relatedness to other MPXV hosts, and the relatedness of MPXV to other poxviruses which are known to infect these hosts. Other major contributing predictors to these five associations were diet composition, habitat landcover, ORF composition in the viral genome, and four-node network features, demonstrating that host ecological traits, molecular virological features, as well as network features, are significantly influencing the predicted host range – underlining the advantage of our multi-perspective approach to investigating virus/host range.

In addition to the European susceptible host hotspots, central China (Sichuan/Gansu provinces) were also highlighted as having a high density of predicted hosts. Of particular note in this region were the Tibetan macaque (*Macaca thibetana*), the Himalayan marmot (*Marmota himalayana*), and the Mongolian gerbil (*Meriones unguiculatus*). Epidemiologically, given the present low prevalence of MPXV in humans in China, spillback is currently of lesser risk in this region. However, it is feasible that the low MPXV prevalence is a result of ongoing SARS-CoV-2 restrictions in the region, and should these restrictions be lifted, MPXV prevalence and therefore spillback risk may increase.

Outside of primates and rodents, our results indicate high probabilities for the domestic cat and dog being susceptible hosts of MPXV (both at >0.88). MPXV infection had not been confirmed for either of these domestic animals at the time of data analysis, however, our data support the current assumption that infection of these animals is feasible, and health organisations’ advice to avoid contact if infected^15^. Since data analysis, the domestic dog has been shown to be susceptible^7^, adding weight to this and confidence in our pipeline’s predictive capacity.

With regards to other wild animals as potential reservoir populations, many close human-associating scavengers, particularly in North America were identified. These include the striped skunk (*Mephitis mephitis*), Virginia opossum (*Didelphis virginiana*), and the common raccoon (*Procyon lotor*), are predicted to be susceptible to monkeypox, all with probabilities above our more stringent 0.76 threshold. Given the large urban populations of these animals, we also suggest them as surveillance priorities.

### Other mammalian poxviruses

Outside of the Monkeypox virus, we studied 50 other fully sequenced mammalian poxviruses, and identified an underestimation of their currently identified host range of between 2.34- and 4.56-fold on average, across all studied poxviruses.

Geographically, hotspots of large numbers of predicted mammalian poxvirus hosts were identified in Western North America, Europe, south-east Africa and central China. The majority of predicted susceptible hosts came from the orders: rodentia, primates, artiodactyla, and carnivora.

Most mammalian poxviruses have a very limited host range, with 30 out of 50 viruses having only one or two known host species. However, like MPXV, both cowpox virus (CPXV) and the closely related vaccinia virus (VACV) have a very wide observed host ranges, spanning 20 and 19 different mammalian families, respectively. Here, we predict an increase of 2.79-4.64-fold (CPXV), and 3.43-7.43-fold (VACV) of susceptible host species. Interestingly, the majority of these predicted susceptible hosts phylogenetically cluster within their known host range (i.e. within the aforementioned families) – particularly expanding the understudied wild species closely related to their well-studied known livestock hosts. These results therefore indicate that there is a much larger pool of wild reservoir hosts in which these economically important viruses may persist. Diligence and separation of livestock from the susceptible wild animals identified here is advisable during CPXV or VACV outbreaks to prevent virus sharing and limit spread.

The top predictors for both CPXV and VACV association with their predicted hosts included: genus ratio, 4-node network predictors, host habitat landcover, and codon biases in the viral genome. This again highlights the importance of a multi-perspective approach to prediction.

The zoonotic orf virus (ORFV) and bovine papular stomatitis virus (BPSV^10^), have significant economic and food security importance for sheep and goats (ORFV^9^) and cattle (BPSV). Of the predicted new host species identified here, for ORFV, 25 (74%), and for BPSV, 35 (76%) were bovids, the majority of which are wild or semi-domesticated. This highlights the underestimated importance of separating livestock from wild bovids, especially during ORFV and BPSV outbreaks.

### Avian poxviruses

Unlike mammalian poxvirus hosts, which span much of the world with a relatively even distribution, the observed and predicted hosts of avian poxvirus are predominantly focused across the palearctic and mainland North America. There was a similar degree of predicted underestimation of host range as for mammalian poxviruses, 1.96-to 4.02-fold on average, across the 12 avian poxviruses studied here.

Of these 12 avian poxviruses, eight have only a single observed host. Amongst those with larger host ranges, canarypox virus (CNPV) and fowlpox virus (FWPV) showed an increase in potential host ranges of 5.1-to 7-fold (CNPV) and 2.72-to 3.44-fold (FWPV).

Unlike for mammalian poxviruses, the most informative predictors for CNPV and FWPV hosts are predominantly a range of phylogenetic, taxonomic and network features. A small to marginal contribution to predictions is made by virus ORF overlap (CNPV) and codon biases (FWPV). Whilst the full range of perspectives did not contribute significantly to the predictions for avian poxvirus hosts, this nonetheless highlights that prediction models focusing on only virus or host perspectives would be significantly less effective than the holistic approach taken here, which enables our pipeline to select the most informative predictors from the full range or any subset of perspectives.

### Concluding remarks

The work presented here shows that there is a significant underestimation in the number of wild animals which could potentially be susceptible to poxviruses, including MPXV. Of most concern is the number of potential MPXV hosts in Europe, the epicentre of the current outbreak, and the clear routes for these species to become infected via contact with contaminated material. The information provided here will enable more focused surveillance, inform policymakers to minimise the highlighted hosts from coming into contact with contaminated material, and potentially help avoid the virus from becoming endemic in new regions.

## Materials and Methods

### Input data

The following input data were used by our model, see Supplementary Note 1 for detailed selection, inclusion, and exclusion criteria:

- **Poxviruses:** Fully sequenced poxviruses obtained from GenBank: 525 sequences of 63 poxviruses (of which 60 had species-level observed hosts).
- **Poxvirus-host associations:** 362 associations between 257 animal species and 60 poxviruses. Data were obtained from ENHanCEd Infectious Diseases Database^16^ (EID2 - https://eid2.liverpool.ac.uk/ - version from May 2022) and pathogen-host associations datasets^17–20^, and supplemented with targeted literature searches (Supplementary Dataset 4). All associations were manually verified for accuracy.
- **Mammalian and avian hosts:** Avian and mammalian species associated with at least one virus (not limited to poxviruses), and a full set of predictors (Supplementary Table 4, Supplementary Note 3) were included (n=2,542). Data were obtained from EID2, pathogen-host associations datasets, and literature searches (Supplementary Dataset 4); and all were manually verified for accuracy.

### Features

We engineered features from virus, host, and network categories (**bold**), grouped them into categories (<u>underlined</u>). For full details of number of features, description and relevance to virus-host interaction see Supplementary Notes 2-4.

#### Virus

<u>Genome</u>: length and GC content. <u>Biases</u>: nucleotide, codon, and amino acid group biases. <u>Secondary structure</u> of predicted proteins: C, E and H coverage. Open reading frames (<u>ORFs)</u>: composition, genome coverage and proportion of total ORF length with overlaps (Supplementary Table 3).

#### Host

<u>Phylogeny/taxonomy</u> Genus and family ratios (ratio of viruses in the game genus, shared with hosts of the same genus/family), phylogenetic distance to known hosts and evolutionary distinctiveness. <u>Host traits</u>: life history traits, diet and, habitats. <u>Geospatial</u>: landcover, climate, size of geographical range, and livestock/poultry headcount within (Supplementary Table 4).

#### Network

<u>3-node motifs</u>; <u>4-node motifs</u> Presents global view of virus sharing amongst hosts, split into multiple motifs or confirmations of possible network structures (summarised in Figure 5, also see Supplementary Table 6).

**Figure 5.**
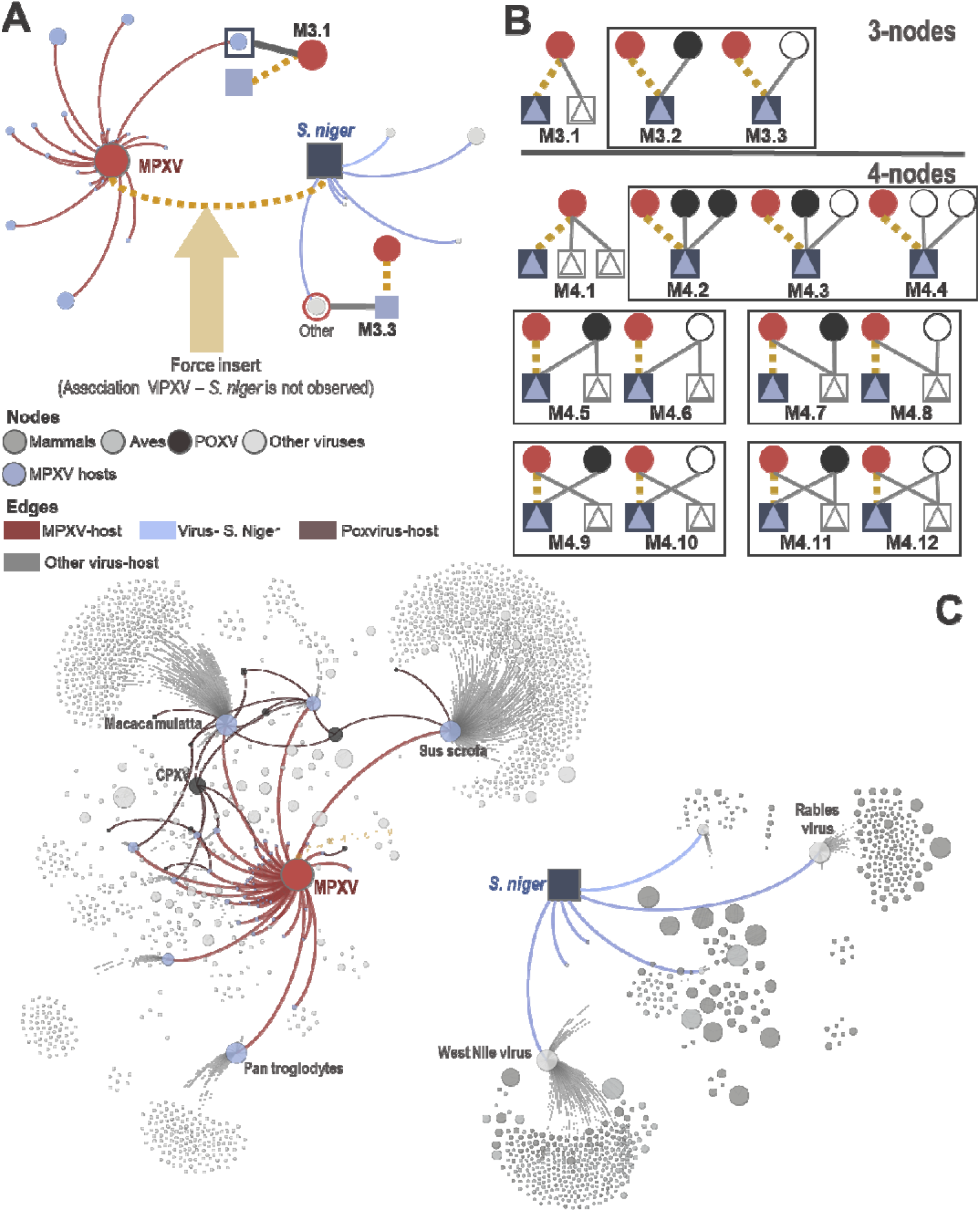
Network features. **Panel A – Potential Motifs in virus-host networks**. The association MPXV-*Sciurus niger* is forced inserted into the network comprising all avian and mammalian viruses and their observed hosts prior to counting motifs (3 and 4-node subgraphs). **Panel B –** 3 and 4-node potential motifs in our virus-host bipartite network. Circles represent viruses and squares with triangles in them represent included animals. Red circles represent the focal virus (v), and blue focal animal species (mammal or aves) of the association for which the motifs are being counted (dashed yellow line). Dark circles represent poxviruses, white circles represent other viruses, white squares with triangles represent hosts. Counts of M3.1 and M4.1 were used directly as features in our models, counts of M3.2, M4.2, M4.3, M4.5, M4.9, and M4.11 were normalised by dividing on total number of motifs in the surrounding box prior to inclusion as features, remainder motifs were not included as features. **Panel C – Motif Space**. Network represents union of 2-step ego networks of MPXV (red circle) and *S. niger* (blue square). Nodes are coloured by category: hosts of MPXV, mammals, aves, poxviruses, and other viruses. Size of nodes is adjusted to represent overall number of hosts or viruses with known associations to the node. Red edges represent nodes reachable from the virus (MPXV) in 1 step (observed hosts). Blue edges represent nodes reachable from the host (*S. niger*) with in one step (known viruses). Dark red edges represent poxvirus-host associations. Light grey edges represent other virus-host associations. Humans were excluded from this network. Motifs are calculated in the union of 1 and 2-step ego networks of both poxvirus and animal.

## Research effort

We calculated research effort into viruses as the total number of sequences and publications of each virus as indexed by EID2^16^. For animal species, we quantified this effort as the total number of sequences and publications of each species, as well as sequences for which the animal species was the host organism, also as indexed by EID2 (Supplementary Note 5).

We then transformed the compound research effort into both virus and animal species into instance weight (Supplementary equations: S.5 and S.6), which enabled us to incorporate research effort directly into the training phase of our LightGBM models. Simply put: observed associations of understudied poxviruses/animals were given slightly more importance than those of over-studied species. Whereas negative (hitherto unobserved) associations of over-studied viruses/animals were given significantly more importance that those of understudied poxviruses/animals. For full details and equations, see Supplementary Note 5.

### Class imbalance

The proportion of observed associations between our poxviruses and their hosts, given all possible associations between our selected poxviruses and included animal species (n= 87,532), was only 0.414%. This considerable imbalance resulted in poor performance of models trained without class balancing (Supplementary Note 5). Due to the small number of observed associations (n=362), we elected to correct for class imbalance using a range of over-sampling and hybrid methods, rather than strict under-sampling. Correction was performed by combining the following techniques: SMOTE (50%), SMOTE (25%) SMOTE-ENN(50%) and SMOTE-ENN (25%), to create a simple averaging ensemble (probability of ensemble = mean probability of constituent models). Supplementary Note 6 lists full details.

### Model training, assessment, and ensemble construction

we performed 100 iterations of model optimisation and training. In each iteration, we split the set of all possible associations (n= 87,532) into training (80%), optimisation (10%), and test (10%) sets. We applied 16 class-balancing to the training set only, and we performed Bayesian (model-based) optimisation of a lightGBM model for each resulting training set against the validation set to tune the hyperparameters. Finally, Performance of tuned models was assessed against the held-out test set. Supplementary Notes 7-8 provide full details.

### Threshold selection

We selected two thresholds to articulate our predictions: >0.50, and ≥0.7602445 (referred to as ≥0.76). 0.5 is the standard threshold. It enhances the ability of our ensembles to detect known associations (higher sensitivity and specificity). Whereas ≥0.76 is calculated such that 90% of observed positives (and detected at threshold > 0.5) are also captured, while reducing number of unknown associations predicted by our models (higher F1-score and higher positive predictive value (PPV/precision), lower sensitivity and specificity).

### Feature importance

We quantified the contribution each of our features (n=216) made on final predictions using SHAP (SHapley Additive exPlanations) values^21^. SHAP values enabled us to measure the extent by which each of those features shaped the probability produced by each constituent model for each possible poxvirus-host association. This in turn enabled us to quantify feature importance at the level of individual viruses or animal species (as well as broader categories), as the mean of absolute SHAP values across all associations for each virus/animal (or category), and all constituent models (n=25×4).

### Data and materials availability

Data and code reported in this paper are available at dx.doi.org/10.6084/m9.figshare.20485332.

## Acknowledgments

MSCB, MB and MW acknowledge BBSRC and NERC grants BB/W00402X/1 and NE/W002302/1 which funded this research.

