## Supplementary materials for "Monkeypox virus shows potential to infect a diverse range of native animal species across Europe, indicating high risk of becoming endemic in the region"

*Supplementary Information*

Marcus SC Blagrove*[1], Jack Pilgrim[1], Aurelia Kotsiri[1], Melody Hui[1], Matthew Baylis[1], Maya Wardeh*[1,2]

*1) Institute of Infection, Veterinary and Ecological Sciences, University of Liverpool, Liverpool, UK*

*2) Department of Mathematical Sciences, University of Liverpool, Liverpool, UK*

### Supplementary Note 1 – Virus-mammal associations data

**Viral genomic data.** Complete sequences of poxviruses were downloaded from Genbank(1). Sequences labelled with the terms: ‘vaccine’, ‘vector’, ‘construct’, or ‘recombinant’ were removed from the analyses. The number of ambiguous bases was then computed for each sequence, sequences with more than 1,024 permutations when ambiguous bases are resolved were removed from the analyses. We further removed viruses that are known to infect non-mammalian and non-avian hosts (e.g., Entomopoxvirinae, Salmonpoxvirus, Crocodilepox virus, Teiidae poxvirus 1, Western carp gudgeon poxvirus). This resulted in 525 sequences of 63 poxviruses, of which, six viruses were unclassified Poxviridae (two mammalian and four avian poxviruses).

**poxvirus-host associations.** Associations between the above 63 poxviruses were compiled from the ENHanCEd Infectious Diseases Database(2) EID2 (https://eid2.liverpool.ac.uk/ - version from May 2022 and pathogen-host associations datasets(3–6), and supplemented those data with targeted literature searches (Supplementary Dataset 4). This resulted in 374 associations between 267 animal species, and 61 poxviruses (two poxviruses did not have species-level hosts in our dataset). All associations were verified manually for accuracy and to remove erroneous interactions.

We elected to train our pipelines with all verified poxvirus-host associations regardless of whether they were obtained by sequence and publications (24.1%), from sequences only (25.1%) or publications only (50.8% - isolation/PCR or serology). Serology is the primary means of determining a prior infection in an individual, whilst PCR, sequencing, isolation, etc. are used to detect a current infection (serology cannot usually identify an early-stage infection, and PCR/sequencing/isolation cannot detect a prior infection). The difference between these diagnostic procedures is therefore methodological and temporal, it does not represent a biological difference between host/virus associations and therefore we see no biological merit in separating the data by “evidence class” or other data curation distinction.

**Selection of potential mammalian and avian hosts of poxviruses.** Combining data from EID2(2) (https://eid2.liverpool.ac.uk/ - version from May 2022), pathogen-host associations datasets(3–7), and literature searches (Supplementary Dataset 4), and following manual verification for accuracy, we identified avian and mammalian species associated with at least one known virus (not limited to poxviruses) (n=2,542). Those animals were included in our models to enable us to generate predictions for hosts which have not yet been associated with any of our 63 poxviruses. We removed any species for which we could not compile a full set of predictors (Table 2, Supplementary Note 3), this resulted in total of 2,484 animal species for which we generated predictions. Additionally, removal of those species resulted in in 362 associations between 257 animal species and 60 poxviruses, which formed our minority positive class (Supplementary Notes 6-8).

**Supplementary Table 1 – Poxviruses included in the study.** Virus classification followed NCBI taxonomy(8). Values in bracket represent standard deviation from the mean.

| | Genus | Poxviruses | Host species | Average host range per virus |
|---|---|---|---|---|
| **Mammalian poxviruses** | **Capripoxvirus** | 3 | 23 | 9 (±9.64) |
| | **Centapoxvirus** | 3 | 2 | 1 (±0) |
| | **Cervidpoxvirus** | 3 | 4 | 1.67 (±1.15) |
| | **Leporipoxvirus** | 2 | 10 | 6 (±2.83) |
| | **Macropopoxvirus** | 2 | 2 | 1 (±0) |
| | **Molluscipoxvirus** | 4 | 3 | 1.25 (±0.5) |
| | **Mustelpoxvirus** | 1 | 1 | 1 (±0) |
| | **Orthopoxvirus** | 18 | 151 | 10.9 (±20.1) |
| | **Oryzopoxvirus** | 2 | 1 | 0.5 (±0.707) |
| | **Parapoxvirus** | 4 | 28 | 9.25 (±7.93) |
| | **Pteropopoxvirus** | 1 | 1 | 1 (±0) |
| | **Sciuripoxvirus** | 1 | 2 | 2 (±0) |
| | **Suipoxvirus** | 1 | 1 | 1 (±0) |
| | **Vespertilionpoxvirus** | 1 | 1 | 1 (±0) |
| | **Yatapoxvirus** | 3 | 10 | 5.67 (±4.04) |
| | **Unclassified poxviruses** | 2 | 2 | 1 (±0) |
| **Avian poxviruses** | **Avipoxvirus** | 8 | 39 | 5.75 (±9.50) |
| | **Unclassified poxviruses** | 4 | 2 | 0.5 (±0.577) |

**Supplementary Table 2 – Domesticated mammalian and avian species included in this study.**

| | Order | Species | |
|---|---|---|---|
| **Mammals** | ***Artiodactyla*** | 15 | *Bison bonasus, Bos frontalis, Bos grunniens, Bos indicus, Bos javanicus, Bos taurus, Bubalus bubalis, Bubalus carabanensis, Capra hircus, Ovis aries, Camelus bactrianus, Camelus dromedaries, Lama glama, Vicugna pacos,* and *Sus scrofa* |
| | ***Carnivora*** | 2 | *Canis lupus familiaris* and *Felis catus.* |
| | ***Perissodactyla*** | 2 | *Equus asinus* and *Equus caballus* |
| | ***Rodentia*** | 5 | *Cavia porcellus, Mesocricetus auratus, Mus musculus, Rattus norvegicus,* and *Rattus rattus.* |
| | ***Lagomorpha*** | 1 | *Oryctolagus cuniculus* |
| **Aves** | ***Anseriformes*** | 4 | *Anas platyrhynchos, Anser anser, Anser cygnoides* and *Cairina moschata* |
| | ***Columbiformes*** | 1 | *Columba livia* |
| | ***Galliformes*** | 3 | *Gallus gallus, Meleagris gallopavo,* and *Numida meleagris* |

**Supplementary Note 2– Poxvirus features**

**Supplementary Table 3 – Viral features groups**. Features were first calculated from sequences and then averaged for each poxvirus. All included viral features are continuous. Detailed methods for features included in each group are listed below.

| Category | Group | Features | Relevance |
|---|---|---|---|
| **Genome** | **Length** | 1 | Affects available space for protein encoding, number of mutations/genome/replication, and size of virion(9). |
| | **GC content** | 1 | GC content affects thermal (and other stressor) stability of genome (and resultant RNA) and regions within them(10). |
| **Biases** | **Nucleotide bias** | 4 | Nucleotide bias can result in genome-wide biases, amino acid composition, and function(11). |
| | **Relative Synonymous Codon Usage (RSCU).** | 64 | Codon biases in viral genome affect translation in host cell. Similarities/differences affect tRNA recruitment, and production speed(12). Can also affect virulence in host(13). |
| | **Amino acids bias** | 19 | Amino acid categories affect protein structure (e.g. proline), pH, solubility, etc. and hence proteins' properties in cellular or extracellular environments(14). |
| **Secondary structure** | **C coverage** | 10 | Secondary structure of proteins is important for final conformation and interaction with host machinery, as well as stability of protein to heat, pH, oxidation etc(15). Similarities in secondary structure can also indicate protein relatedness not identifiable by sequence(16). |
| | **E coverage** | | |
| | **H coverage** | | |
| **ORFs** | **Composition** | 4 | Proportion of genome utilised by ORFs, the sizes of the ORFs, and the proportion of overlapping ORFs (in different frames between) describes genome utilisation |

| | | | |
|---|---|---|---|
| | **Coverage** | 2 | for protein production, genome density, and constraints on AA sequence (from overlap). |
| | **ORF Overlap** | 7 | |

**Sequence cleaning and ORF generation.** Sequences with ambiguous bases that could be resolved in fewer than or equal to 1,024 permutations (n=525) were expanded to resolve ambiguity using Disambiguate function in the R package Decipher(17). This resulted in 9,304 cleaned sequences. ORFs (non-overlapping set containing longest ORF) were then predicted for each sequence and its reverse complement using predORF function in the R package systemPipeR (parameters were set to n='all' and longest_disjoint=TRUE to subset to non-overlapping ORF set containing longest ORF. Predicted ORFs of length <123 were dropped from the analyses.

**Genome wide features.** GC content, Nucleotide biases (proportion of each nucleotide in the sequence) and genome length were computed for each of the cleaned sequences. Values were then averaged per each poxvirus to generate final features.

**ORF Composition.** We computed the following feature to express sense and asense ORF composition for each sequence:

1. Sense bias = number of ORFs (length≥123) predicted from each sequence ($n_{sense}$) divided by sequence length.
2. Asense bias = number of ORFs (length≥123) predicted from the reverse complement of each sequence ($n_{asense}$) divided by sequence length.
3. Sense Probability = $\frac{n_{sense}}{n_{sense} + n_{asense}}$
4. Asense/Sense probability = $\frac{n_{asense}}{n_{sense}}$

**ORF coverage & overlap.** For each sequence, we computed a coverage vector (length = sequence length, initialised with 0s). For each (non-overlapping) predicted $ORF_i$, coverage vector is updated such that:

$$Coverage\left(start_{ORF_i}, end_{ORF_i}\right) = inframe2end_{ORF_i} \qquad \textbf{(S.1)}$$

Where $inframe2end_{ORF_i}$ is frame of identified ORF/CDS relative to 3' end of query sequence. For ORFs predicted from each sequence (sense), $inframe2end_{ORF_i} \in \{1,2,3\}$, where value 1 stands for in-frame with downstream ORF, whereas 2 or 3 indicates a shift of one or two bases, respectively. For ORFs predicted from reverse complements (asense), $inframe2end_{ORF_i} \in \{4,5,6\}$, where value 4 stands for in-frame with downstream ORF, whereas 5 or 6 indicates a shift of one or two bases, respectively.

1. Sense coverage = proportion of elements in coverage with value ∈ {1,2,3},
2. Asense coverage = proportion of elements in coverage with value ∈ {4,5,6},

Additionally, we expressed ORF overlap in 7 features ($T_0$ to $T_6$), whereby, for each possible frame value $k$ (0 = no predicted ORFs, 1 = in-frame with downstream ORF, 2 or 3 a shift of one or two bases, 5 = in-frame with downstream ORF (reverse complement), 5 or 6, a shift of one or two bases), $T_k$ = proportion of elements in coverage = $k$.

**Relative Synonymous Codon Usage (RSCU).** Frequency of each codon (n = 64) was calculated for each predicted ORF. For each sequence, frequencies were then summed for all ORFs obtained from a said sequence. RSCU was then computed for each codon (including stop codons)(18) as follows: let $n_i$ be the number of codons synonymous for amino acid $AA_i$ and $C_{ij}$ the frequency of the $j^{th}$ codon encoding for $AA_i$ then:

$$RSCU(C_{ij}) = \frac{C_{ij}}{\frac{1}{n_i} \times \sum_j^{n_i} C_{ij}} \qquad \textbf{(S.2)}$$

**Amino Acids biases.** Frequency for each Amino acid was computed for each predicted ORF, and then summed across all ORFs obtained from the same sequence. Amino acids were categorised into 19 overlapping binary categories expressing:

1. Hydropathy: neutral, hydrophilic, hydrophobic.
2. Volume: very large, large, medium, small, very small.
3. Charge: negative, positive.
4. Polar.
5. Hydrogen donor or acceptor.
6. Chemical: acidic, aliphatic, amide, aromatic, basic, hydroxyl, and Sulphur.

Bias for each amino acid category was then computed as frequency of all amino acids encoding for each category in a sequence, divided by the total number of amino acids in the sequence.

**Secondary structure.** Full length poxvirus genomes were aligned using the 'fftns' algorithm in MAFFT v7.490(19). Gblocks v0.91b(20) was then used to exclude areas of poor alignment using '-b5=h' to partially allow for gap positions. Following alignment of poxvirus sequences (above), we predicted the secondary structure for each sequence using PredictHEC function in the R package Decipher(17). We then computed for each 20% of the genome length (both sense and anti-sense) the coverage (number of times a structure was predicted). Thus we generated 30 features for each genome representing coverage for each of the three possible structures—Helix (H, 10 features), Beta-Sheet (E, 10 features) or Coil (C, 10 features).

### Supplementary Note 3 – Mammalian and avian features

**Supplementary Table 4 – Host features. Supplementary Note 3 lists full details for all host features.**
Detailed methods for each are feature group are listed below.

| Category | Group | Feature(s) | Data type | Relevance |
|---|---|---|---|---|
| **Phylogeny/ taxonomy** | **Genus ratio** | Genus ratio | Continuous | For each individual poxvirus/host prediction: Number of poxviruses in the same genus (as the poxvirus in question) known to affect other hosts in the same genus/family as the potential host in question (divided by total possible number of associations for that genus/genus or genus/family combination) |
| | **Family ratio** | Family ratio | | |
| | **Phylogeny** | Phylogenetic distance to known hosts | | Indicated, for each poxvirus/animal combination (n= 87,532), whether the animal is phylogenetically close or distant from the poxvirus known host range |
| | | Evolutionary distinctiveness (ED) | | ED expresses the degree of isolation or connectivity of a species in its phylogenetic tree(21), and has been shown to correlate negatively with pathogen species richness(22). |
| **Host traits** | **Life history traits** | Body mass (grams) | | Host life history traits and physiological phenotypes correlate with and can explain reservoir competence(23,24). |
| | | Gestation period (days) | integer | |
| | | Clutch size | | |
| | | Volant | binary | |
| | | Activity cycle (4 categories: diurnal, nocturnal, cathemeral, and crepuscular) | | |
| | **Diet** | Proportional use of 10 diet categories(25). | Continuous | Diet has been shown to affect virus competence of mammalian species(26). And shared diets/predation can affect host interaction and hence propensity to share viruses. |
| | | Diet entropy: Shannon entropy of diet proportional use. | | |
| | **Habitats** | Habitat utilisation (15 categories) | Integer | Animals utilising similar habitats might encounter similar viruses, hence increase the chances of being infected with these viruses |
| | | Artificial habitat | binary | Indicating whether species utilise any artificial habitat. |
| | | Habitat breadth | Integer | Indicating the overall breadth of species habitat preference/utilisation.<br>Habitat breadth = sum of total habitat used (range between 1 and 75).<br>Unique habitats = sum of habitat categories used (range between 1 and 15). |
| | | Unique habitats | | |
| **Geospatial** | **Landcover** | Mean and SD in 12 categories | Continuous | Landcover type affects distribution of hosts(27), more overlapping of landcover usage between hosts could indicate more direct or indirect interaction and sharing of viruses. |
| | **Livestock and poultry headcount** | Bovid livestock species/pigs/horses/poultry/pig and poultry | | Livestock farming is linked to emergence and cross-species transmission of many viruses(28). |
| | **Climate** | Mean and SD for temperature/ Diurnal temperature range/ precipitation | | Climate influences many human and domestic animal pathogens(29–31). Climate also affects arthropod vectors, which mechanically transmit ~20spp of poxvirus(32). |
| | **Area** | Geographical range (area size in $km^2$) | | More widely distributed species might be exposed to more viruses. |

**Genus and family ratios**. We computed two features based on the taxonomic relations between each animal species ($h_j$) and known hosts of each poxvirus ($v_i$) as follows:

1. Let $Observed\left(G(v_i), G(h_j)\right)$, be the number of observed associations between viruses in the same genus as $v_i$, and animals in the same genus as $h_j$; and $Possible\ \left(G(v_i), G(h_j)\right)$, be the number of all

possible associations (observed or not) between viruses in the same genus as $v_i$, and animals in the same genus as $h_j$; then:

$$Genus\ Ratio\ (v_i, h_j) = \frac{Observed\ \left(G(v_i), G(h_j)\right)}{Possible\ \left(G(v_i), G(h_j)\right)} \quad \textbf{(S.3)}$$

2. Let $Observed\ \left(G(v_i), F(h_j)\right)$, be the number of observed associations between viruses in the same genus as $v_i$, and animals in the same family as $h_j$; and $Possible\ \left(G(v_i), F(h_j)\right)$, be the number of all possible associations (observed or not) between viruses in the same genus as $v_i$, and animals in the same family as $h_j$; then:

$$Family\ Ratio\ (v_i, h_j) = \frac{Observed\ \left(G(v_i), F(h_j)\right)}{Possible\ \left(G(v_i), F(h_j)\right)} \quad \textbf{(S.4)}$$

We repeated the calculations per each iteration of our training and model selection process (n=100), such that "observed hosts" were determined by the number of poxvirus -host associations available for each of training, optimisation and validation sets as follows:

1. Training: only poxvirus-host associations within the training set (~80% of total) were used.
2. Optimisation: poxvirus-host associations within both training and optimisation sets were used (~90%).
3. Validation: all available poxvirus-host associations (n=362) were used.

This measure indicated, for each poxvirus/animal combination (n= 87,532), whether the animal species is phylogenetically close or distant from the poxvirus known host range.

**Phylogeny**. Phylogenetic classness have been found to drive spill-over and sharing of viruses(4,33,34). We obtained recent phylogenetic trees for mammals(35), and avian(36) species from vertLife (http://vertlife.org/). We built majority-rule (with branch length) consensus tree for mammals (and similarly for avian) species, by drawing 1,000 trees from the downloaded phylogenies, and applying consensus.tree and consensus.edges from the R package phytool as follows.

Mean phylogenetic distance to known hosts: We utilised the above consensus trees to calculate pairwise phylogenetic distance between each mammal-mammal (n=2,217,121) and aves-aves (n=990,025) pair. We then computed the mean distance between each mammalian species and the known hosts of each of our mammalian poxviruses (n=51), and for each avian species and the known hosts of each of our avian poxviruses. (n=12). Calculations were conducted per each iteration of the training processes (n=100) as per genus and family ratio calculations.

Evolutionary distinctiveness: We computed the evolutionary distinctiveness for each animal species (n= 2,484) using fair proportion(37), as implemented in the R package picante(38). Evolutionary distinctiveness quantifies how isolated or connected a species is on its phylogenetic tree(21), and has been shown to correlate negatively with pathogen species richness(22).

**Species traits.** We compiled data on morphological, reproductive and life-history traits, diet and habitat of our avian and mammalian species from online databases and literature(25,39–46). Supplementary Table 4 lists the traits we have selected the following traits for their known correlation with virus-host associations, their wide availability, and their applicability to both avian and mammals.

**Habitat utilisation:** Instead of using binary (yes/no) indicators of whether an animal use specific habitat (e.g., forest), we incorporated habitat utilisation(41) as multiple numerical indicators (following IUCN classification scheme - https://www.iucnredlist.org/resources/habitat-classification-scheme) of whether a species uses one or more of 102 natural and artificial habitats in 15 categories. For instance, animals known to utilise both temperate and subtropical/tropical forests would have forest = 2, instead of forest = yes.

We merged underutilised habitats in each category to one subcategory (other or subarctic/subantarctic). This resulted in 75 habitats/subcategories as follows: forest = 8, savanna =2, shrubland = 7, grassland = 6, wetlands = 9, rocky areas = 1, caves & subterranean = 2, desert = 3, marine neritic=9, marine oceanic = 3, marine intertidal = 7, marine coastal/supratidal = 4, artificial terrestrial = 6, artificial aquatic = 6, and other=1.

**Geospatial features.** We obtained species-presence maps for majority of our avian species from BirdLife(47), and mammalian species from IUCN(41). We extrapolated livestock (including horses) species-presence maps from most recent global distribution maps(48). Finally, we inferred presence-maps for three domesticated species - dogs (*Canis lupus familiaris*), cats (*Felis catus)* and guinea pigs (*Cavia porcellus*) from gridded population of the world maps(49), by assuming they co-exist with humans where there is sufficient human populations ($n$>100). We used the same gridded population maps to extrapolate human species-presence map ($n$>0). We supplemented those maps with grids expressing climate(50), landcover (including urbanisation)(51)and distribution of poultry and livestock(48). Supplementary Table 5 lists these sources, their resolution and resulting features.

This enabled us to generate the following geospatial features of our avian and mammalian species (

1. We expressed climate in 6 features: mean and standard deviation (SD) of mean temperature (within species range), mean and SD of precipitation, mean and SD of diurnal temperature range.
    a. Mean temperature: mean of monthly temperatures recorded in the species-presence area, averaged between years: 1900-2010(50).
    b. Precipitation: Sum of monthly precipitation recorded in the species-presence area, averaged between years: 1900-2010(50).
    c. Diurnal temperature range (DTR): mean of monthly DTR recorded in the species-presence area, averaged between years: 1900-2010(50).
2. We expressed each land cover category (12 in totals – value between 0 and 100% per each grid cell) as mean and SD (24 features in total).
3. We quantified head count of domesticated livestock (including horses) and poultry in the species presence area into the following features:
    - Head count of bovid livestock (including cattle, sheep, goats, and buffaloes)
    - Head count of pigs
    - Head count of horses
    - Head count of poultry (chicken and ducks)
    - Head count of poultry and pigs where they co-exist in the same grid cell.
4. We included geographical area range (in $km^2$) as species with wider areas might be exposed to more viruses.

**Supplementary Table 5 - List of geographical layers integrated within our models.**

| | **Geographical layer** | **Source** | **Resolution** |
|---|---|---|---|
| **Land-cover** | Evergreen/deciduous needle-leaf trees (%) | EarthEnv(51) | 0°0′30″ |
| | Evergreen broad-leaf trees (%) | | |
| | Deciduous broad-leaf trees (%) | | |
| | Mixed/other trees (%) | | |
| | Shrubs (%) | | |
| | Herbaceous vegetation (%) | | |
| | Barren land (%) | | |
| | Managed/Cultivated Vegetation (%) | | |
| | Regularly flooded vegetation (%) | | |
| | Urban (%) | | |
| **Livestock and poultry head count** | Cattle (head count) | Global distribution data (livestock)(48) | 0.0833° |
| | Sheep (head count) | | |
| | Buffalo (head count) | | |
| | Pigs (head count) | | |
| | Horses (head count) | | |
| | Chicken (head count) | | |
| | Duck (head count) | | |
| **Climate** | Mean temperature | CRUTS3(50) | 0°5′ |
| | Precipitation | | |
| | Diurnal temperature range | | |

### Supplementary Note 4 – Network features

We constructed a bipartite network linking 11,198 viruses and 2,484 hosts and comprising 24,445 edges (host-virus associations). This network presented a global view of sharing of viruses amongst mammalian and avian hosts, which enabled us to quantify potential pathways of sharing of poxviruses. We captured these pathways by means of counts of potential motifs(52,53) - simple (3 and 4-node) sub-graphs featuring potential poxvirus-animal species associations (Figure 5-A:C), which enabled integration of network structure directly into our predictive pipelines.

These motifs represent important pathways between poxviruses and their potential hosts, varying from simple generalisations to more complex indirect pathways (Supplementary Table 6).

**Supplementary Table 6 – Network features (counts of potential motifs).**

| Feature | Nodes | Focus | Relevance |
|---|---|---|---|
| $M_{3.1}$ | 3 | poxvirus | Whether the focal poxvirus has a wide or narrow observed host range. |
| $M_{4.1}$ | 4 | | |
| $M_{3.2}$ (normalised = $M_{3.2}$/( $M_{3.2}$+ $M_{3.3}$)) | 3 | animal species | Whether the focal animal species is proportionally exposed to many poxviruses or not. |
| $M_{4.2}$ (normalised = $M_{4.2}$/( $M_{4.2}$+ $M_{4.3}$+ $M_{4.4}$)) | 4 | | |
| $M_{4.3}$ (normalised= $M_{4.3}$/( $M_{4.2}$+ $M_{4.3}$+ $M_{4.4}$)) | 4 | | |
| $M_{4.5}$ (normalised = $M_{4.5}$/( $M_{4.5}$+ $M_{4.6}$)) | 4 | | Whether the focal animal species is proportionally exposed to multi-host poxviruses, given all multi-host viruses known to infect it. |
| $M_{4.7}$ (normalised = $M_{4.7}$/( $M_{4.7}$+ $M_{4.8}$)) | 4 | poxvirus | Whether the focal poxvirus infects hosts that are exposed to other poxviruses, factoring in other viruses that infect these hosts. |
| $M_{4.9}$(normalised= $M_{4.9}$/( $M_{4.9}$+ $M_{4.10}$)) | 4 | association | Proportion of indirect links that the association would form between hosts of other poxviruses, given the observed host range of the focal poxvirus. |
| $M_{4.11}$(normalised= $M_{4.11}$/( $M_{4.11}$+ $M_{4.12}$)) | 4 | | Number of 4-node cliques that might form should the focal association exist. |

### Supplementary Note 5 – Research effort

We incorporated research effort on animal species and poxviruses directly into the training phase of our models. We calculated research effort into viruses as the total number of sequences and publications of each virus ($RE_v$) as indexed by EID2(2). For animal species, we quantified this effort ($RE_h$) as the total number of sequences and publications of each species, as well as sequences for which the animal species was the host organism, also as indexed by EID2. We then transformed the compound research effort into both virus and animal species, for each possible virus-animal association ($v_i h_j$), into association (termed instant) weight in two steps.

Firstly, we computed association weight in given training sets

$$W_{v_i h_j} = 1 - \frac{RE_{v_i}}{\sum_{v \in Poxviruses} RE_v} \times\ 1 - \frac{RE_{h_j}}{\sum_{h \in Animals} RE_h} \quad \textbf{(S.5)}$$

Secondly, adjusted $W_{v_i h_i}$ to take into account whether the association $v_i h_i$ has been observed (or not), and whether the virus and the animals are over- or under-studied as follows: let $avg_V$ be the mean research effort into poxviruses in a given set, $avg_H$ the mean research effort into animal species, and $O$ the set of observed associations in the same set, then:

$$AW_{v_i h_j} = \begin{cases} W_{v_i h_j} + 1.5\ v_i h_j\ \notin O\ \wedge RE_{v_i} \le avg_V \wedge RE_{h_j} \le avg_H \\ W_{v_i h_j} + 3\ v_i h_j\ \in O\ \wedge RE_{v_i} \le avg_V \wedge RE_{h_j} \le avg_H \\ W_{v_i h_j} + 2\ v_i h_j\ \in O\ \wedge RE_{v_i} > avg_V \wedge RE_{h_j} > avg_H \\ W_{v_i h_j}\ otherwse \end{cases} \quad \textbf{(S.6)}$$

In other words, observed associations of understudied poxviruses/animals were given slightly more importance than those of over-studied species. Furthermore, negative (hitherto unobserved) associations of over-studied viruses/animals were given significantly more importance that those of understudied poxviruses/animals.

### Supplementary Note 6 – Class balancing

Each poxvirus in our dataset affected 3.45 animals on average(~0.240% of the 1,436 mammals in our models), and each mammalian host was affected by 4.41 viruses (~0.241% of the 1,833 viruses in our models). This presented considerable imbalance in the distribution of observed (known = positive), and unknown (=negative) virus-host associations in our data. This considerable imbalance resulted in poor performance of models trained without class balancing (Supplementary Figure 1).

Due to the small number of observed associations (n=362), we elected to correct for class imbalance using a range of over-sampling and hybrid methods, rather than strict under-sampling. Supplementary table 5 lists the techniques included in this study. All balancing was applied to training sets prior to training and was performed using R packages: bimba and imbalance (ENN only).

Additionally, we compared performance of our selected techniques with tuning the scale_pos_weight parameter of the underlying lightGBM algorithm (R package lightgbm). This parameter defines the ratio of the negative class to the positive class and allows to assign a configurable (tuneable) weight to the minority class. Supplementary Figure 1 illustrates the performance of models trained with this parameter being the in the tuneable hyperparameter set (Supplementary Note 7).

**Supplementary Table 7 – Class balancing techniques used in this study.** % refers to percent of minority class instances (here positive poxvirus-host association) in the resulting training set.

| Technique | Sampling | Method | % |
|---|---|---|---|
| **SMOTE** | **Over-sampling** | SMOTE (Synthetic Minority Over-Sampling Technique(54)) synthesises new minority class instances (here: poxvirus infects animal) from existing cases using a k-nearest neighbour algorithm. SMOTE then over-samples from the minority instances (original and synthesised) and under-samples from the majority class to create a balanced training set. | 50% |
| **SMOTE (25%)** | | | 25% |
| **BL-SMOTE** | | Borderline-SMOTE(55) uses the number of minority class neighbour to classify each minority class instance into either noise (where number of minority neighbours = 0); or safe (where minority neighbours ratio >50%). Both sets are excluded from the synthesising process, and new instances are generated along the decision boundary between minority and majority classes (unlike SMOTE, which generates minority instances randomly). Borderline-SMOTE has two options, here we adopted option 2, whereby minority classes are oversampled, instead of option 1 which oversamples from both classes. | 50% |
| **BL-SMOTE (25%)** | | | 25% |
| **SL-SMOTE** | | Safe-Level-SMOTE(56) defines “safe level” as the number of majority instances in k nearest neighbours of an instance of the minority class. Where the safe level of an instance is close to 0, the instance is classified as noise, whereas if it is close to k, the instance is considered safe. However, unlike borderline-smote, safe-level-smote synthesises its minority instances in “safe positions” by factoring in the safe level ratio of said instances, thus avoiding using outlier minority instances when generating new instances. | 50% |
| **SL-SMOTE (25%)** | | | 25% |
| **ADASYN** | | Adaptive Synthetic Sampling (ADASYN)(57) adaptively synthesises minority instances in inverse proportion to the density of minority instances in their neighbourhood. In other words, ADASYN generates more instances in the regions of the feature space where the density of minority instances is low, and fewer or none where the density is high. As such, ADASYN differs from SMOTE in that it automatically adjusts number of new instances generated for each minority instance to compensate from skewed distributions, instead of generating the same number of synthetic instances for each original minority instances. | 50% |
| **ADASYN (25%)** | | | 25% |
| **SMOTE (NRAS)** | **Noise reduction & over-sampling hybrid** | Noise Reduction A Priori Synthetic Over-Sampling (NRAS)(58) is applied to “clean” training data prior to synthesising new instances using SMOTE, Borderline-SMOTE, Safe-Level-SMOTE and ADASYN. NRAS removes minority instances that have the proportion of minority examples among their k nearest neighbours below a threshold (default = 50%). Note that in this study we utilised NRAS to clean training data, as implemented in the R package bimba (which has been uncoupled from the SMOTE step in the original implementation(58)). We subsequently applied over-sampling algorithms the cleaned data. | 50% |
| **BL-SMOTE (NRAS)** | | | 50% |
| **SL-SMOTE (NRAS)** | | | 50% |
| **ADASYN (NRAS)** | | | 50% |
| **SMOTE-ENN** | **Over- & under-sampling hybrid** | SMOTE(56) followed by Edited Nearest Neighbours (ENN)(59). ENN labels instances whose class labels differ from the class of at least two of their three nearest neighbours as noise, and removes them from the training set, thus creating a smoother decision surface. | 50% |
| **SMOTE-ENN (25%)** | | | 25% |

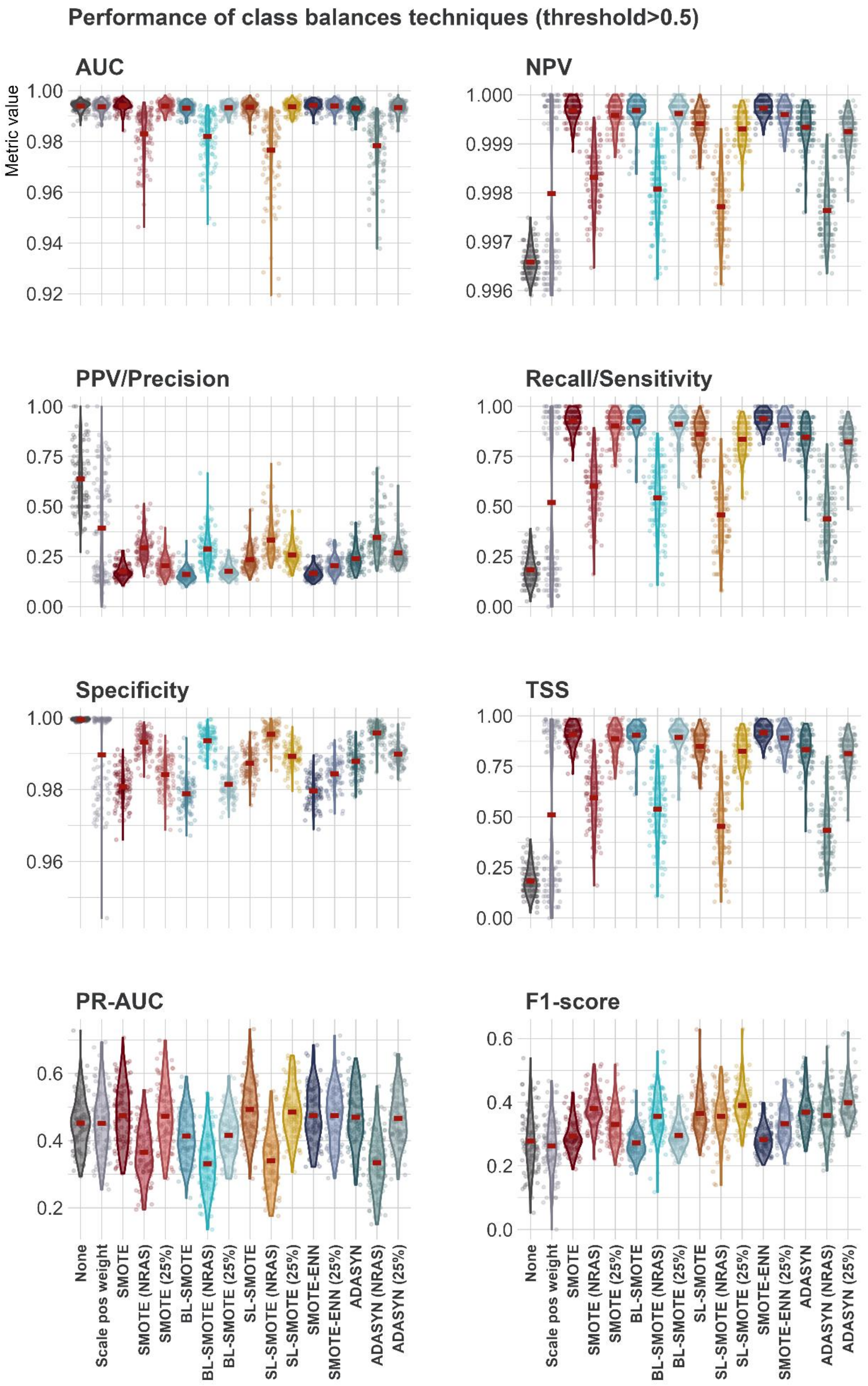


**Supplementary Figure 1 – Performance assessment of class balancing techniques over all held-out test sets (n =100) at >0.5 probability threshold.** Points represent results from individual iterations (n=100). Violin plots show the kernel probability density of the data at different values. Points and violin plots are coloured by class balancing algorithms. Each held-out test-set comprise a 10% stratified random sample of all possible associations (n= 87,532).

**Supplementary Note 7 – Model training, optimisation, and validation**

**LightGBM** (Lightweight Gradient Boosting Machines)(60) is an open-source gradient boosting framework which uses tree-based learning algorithms. Gradient boosting machines build decision trees (weak learners) sequentially. Each tree is built based on the error obtained from previous trees, and predictions are generated based on the sum of all trained tree. Unlike other implementations (e.g., XGBoost) which grow trees in level-wise manner, LightGBM implements a leaf-wise approach to tree growth, which enhances training and prediction speed of LightGBM models.

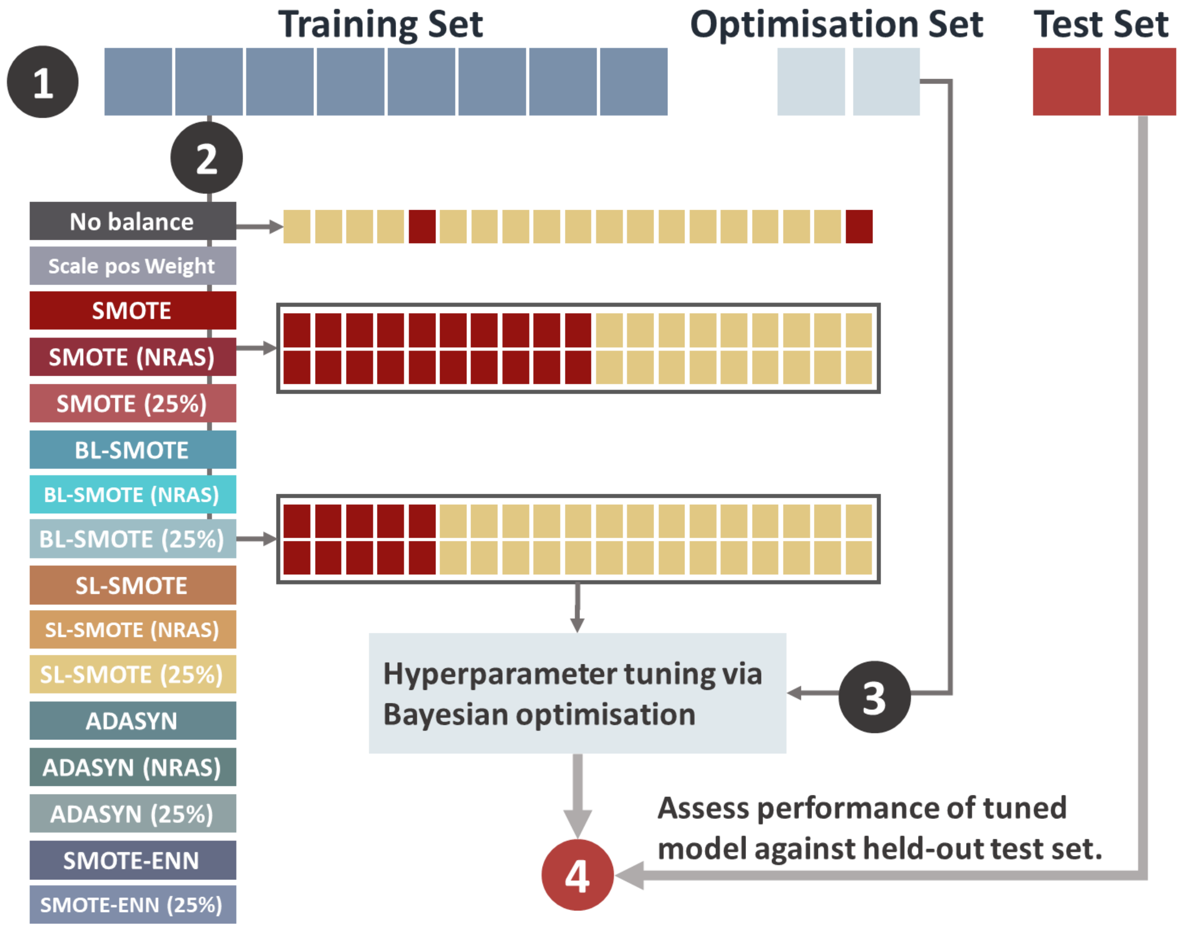


**Supplementary Figure 2 – Training, optimisation and validation of individual model.**

Here, we performed 100 iterations of tuning, training, and validation of LightGBM boosted decision trees models, using the R Packages: mlr3, mlr3extralearners, mlr3tuning, mlrintermbo, and lightgbm, as follows (Supplementary Figure 2):

- **Step 1 -** Per each iteration (n=100), we split the set of all possible associations (n=87,532) between fully sequenced poxviruses (n=63) and animal species (n=2,484) into three sets: training (80%), optimisation (10%) and validation (held-out test, 10%) sets. Sets were draw using stratified random sampling to maintain class ratio (positive=observed/known) in the resulting splits. Interaction derived features (host/genus ratios, phylogenetic distance to known hosts, network features) and training weights were then computed for each iteration.
- **Step 2 –** Each training set was then passed through our balancing pipeline, which generated 15 training sets (original training set (unbalanced), and 14 balanced using the techniques listed in Supplementary Note 6). Integer and binary variables were adjusted post balancing by rounding. Research effort into both viruses and animals was incorporated in the balancing process. Instance weights were then derived for resulting sets from the resulting research efforts.
- **Step 3 –** A LightGBM model with instance weights was then optimised for each of the resulting training sets against the validation set (10%, not balanced, same set was used for all models trained per each iteration). We trained 16 models in total per each iteration:

a. Two models were tuned without balancing classes (original training set): no balance (original training data), scale positive weight (original training data + scale_pos_weight as hyperparameter (tuned in range [1:5000])).
b. 14 models were trained with the balanced/semi-balanced training sets resulting from our balancing pipeline.

We performed Bayesian (model-based) optimisation against the validation set to tune the hyperparameters. We set the tuning stopping to stagnation over 15 rounds, and optimisation was performed using the rpackages: mlr3tuning and mlrintermbo.

- **Step 4** – performance of tuned models was assessed against the held-out test sets (validation sets = 10%). Performance of class balancing techniques (Supplementary Note 5) was compared across the 100 iterations. Predictions from four techniques: SMOTE, SMOTE (25%), SMOTE-ENN and SMOTE-ENN (25%) were then averaged (per iteration) to generated final ensemble.

**Supplementary table 8.** LightGBM hyperparameters tuned per iteration.

| **Parameter** | **Range** | |
|---|---|---|
| num_leaves | [7:4095] | Defines maximum number of leaves per weak learner (tree). Larger values increase accuracy on the training set but might lead to overfitting. |
| max_depth | [2:63] | Controls the maximum depth of each tree within the model, it is trained in tandem with num_leaves. Larger values increase accuracy on the training set but might lead to overfitting. |
| min_data_in_leaf | [200:10000] | Defines the minimum number of instances that must be contained in a leaf to be added to the tree. It controls for overfitting, so that the model does not become too specific. |
| Lambda_l1 | [0:100] | Regularisation parameters used to control overfitting. |
| Lambda_l2 | [0:100] | |
| min_gain_to_split | [0:15] | Regularisation parameter. When adding a new tree node, LightGBM chooses the split point that has the largest gain. Simply put, gain is the reduction in training loss that results from adding a split point. Larger values decrease training time. |
| min_sum_hessian_in_leaf | [0.01: \|train\|/1000] | The sum of hessians of the instances contained in the leaf. Hessian of a data point is the second order derivative of the loss function evaluated for each instance. It takes instance weights into consideration, as hessians are multiplied by those weights prior to passing them to the weak learner (tree). Tuning this hyperparameters allows weak learners to perform a more flexible split for those data instances with low confidence, and less flexible split for those with high confidence, thus providing an adaptive regularisation. |
| bagging_fraction | [0.4:1] | Controls the size of sample used in constructing each weak learner. For instance, a value = 0.7, means that each tree will be constructed using 70% of training data, randomly sampled without replacement. Lower values decrease training time. |
| feature_fraction | [0.4:1] | LightGBM randomly selects a subset of features per each tree (weak learner), feature_fraction defines % of feature included in each tree. For instance, when set to 0.4 LightGBM will sample 40% of features to be included in training each tree. This hyperparameter has two uses: speeding up training and avoiding overfitting. |

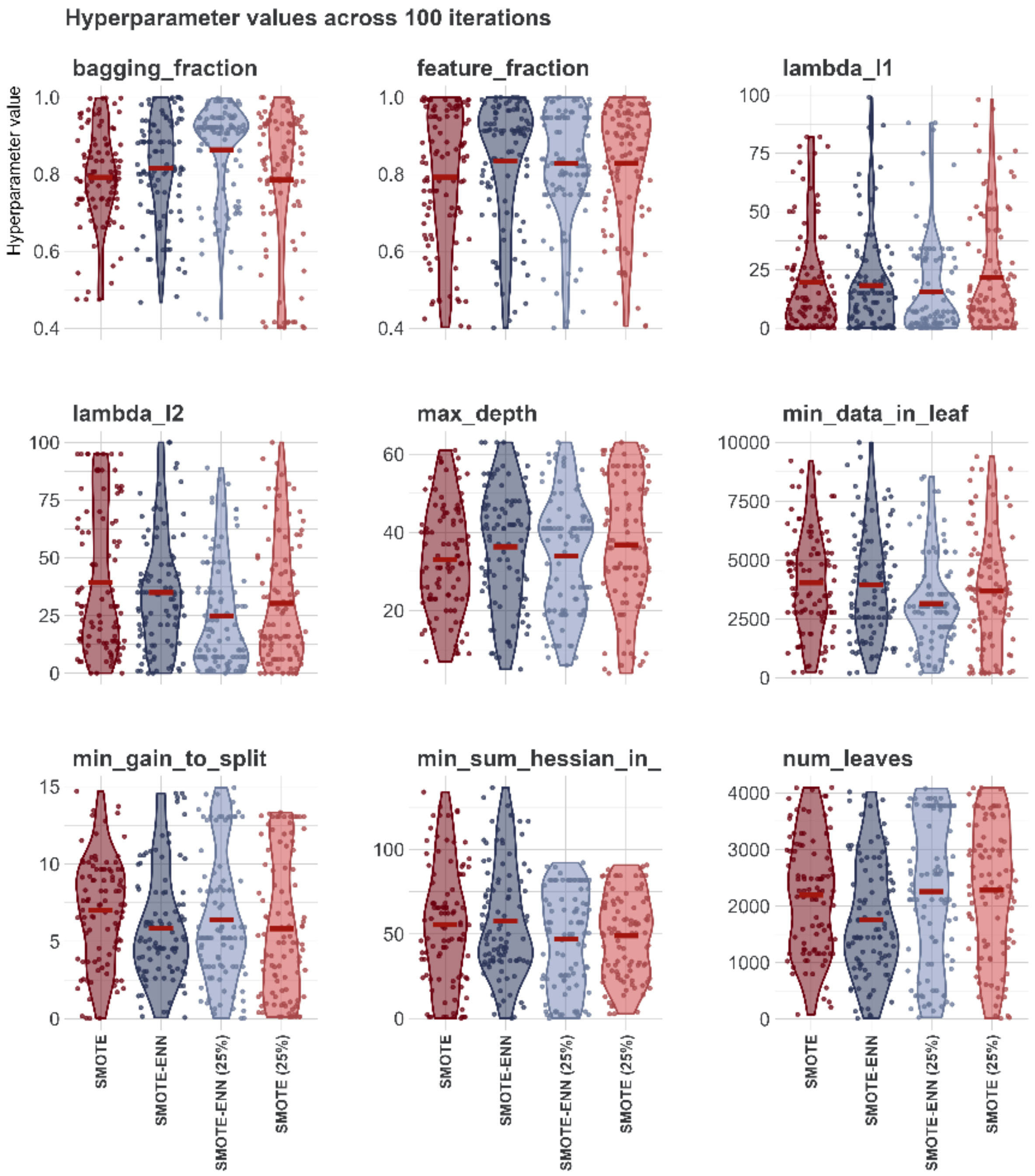


**Supplementary Figure 3 – LightGBM Hyperparameters.** Points represent results from individual iterations (n=100). Violin plots show the kernel probability density of the data at different values. Points and violin plots are coloured by class balancing algorithm: SMOTE, SMOTE (25%), SMOTE-ENN, and SMOTE- ENN (25%).

### Supplementary Note 8 – Performance metrics

**Supplementary Table 9 - Measures utilised to assess the performance of our ensembles and their constituent models.** Bolded measures were used in ranking and selecting top 25 performing ensembles.

| **Confusion matrix** | Detected | |
|---|---|---|
| Predicted | 1 | 0 |
| 1 | TP (true positives) | FP (False positives) |
| 0 | FN (False negatives) | TP (true negatives) |

| **Measure** | **Formula** | **Meaning** |
|---|---|---|
| Sensitivity (recall) – True Positive Rate (TPR) | $\frac{\text{TP}}{\text{TP} + \text{FN}}$ | Sensitivity is the percentage of actual positives (observed associations) that were correctly predicted. It indicates the percentage of 1s that was covered by the model. |
| False Positive Rate (FPR) | $\frac{\text{FP}}{\text{FP} + \text{TN}}$ | Percentage of false positive predictions of the model. |
| Specificity | $\frac{TN}{\text{TN} + \text{FP}}$ | Specificity is the percentage of negatives (here unknown associations, not necessarily true negative) that were correctly predicted |
| Precision (PPV (Positive Predictive Value)) | $\frac{\text{TP}}{\text{TP} + \text{FP}}$ | Percentage of accurate positive predictions of the model. |
| NPV (Negative Predictive Value)) | $\frac{TN}{\text{TN} + \text{FN}}$ | Percentage of accurate negative predictions of the model. |
| **AUC** | **Area Under the ROC Curve** | Threshold-independent measure of model predictive performance that is commonly used as a validation metric for host-pathogen predictive models(61,62). AUC favours both classes (negative/positive) equally. AUC captures how well the model separates the positive and negative examples and is calculated based on the TPR and FPR values. |
| TSS | $\text{Sensitivity} + \text{Specificity} - 1$ | Use of AUC has been criticised for its insensitivity to absolute predicted probability and its inclusion of a priori untenable prediction (63,64), we also calculated the True Skill Statistic (TSS)(65). |
| **PR-AUC** | **Area Under the (precision-recall) Curve** | Threshold-independent measure, widely used with uneven class distribution. PR-AUC favours the positive (minority) class, and is calculated based on the TPR and PPV values. |
| F1-score | $2 \times \frac{\text{Precision} \times \text{Recall}}{\text{Precision} + \text{Recall}}$ | Captures the harmonic mean of the precision and recall. F1-score (F-score for short) is often used with uneven class distribution. |

### Supplementary Note 9 – Methodological limitations

We acknowledge certain limitations in our methodology, primarily pertaining to current incomplete data sets:

- We were limited to using only fully sequenced poxviruses, as the full sequence was required to generate genomic features. The same applies for host species — we could only include hosts for which phylogenetic, ecological, and geospatial data were available. This unavoidably limited the virus and host range used in the study.
- For the vast majority of observed poxvirus–host associations it is unknown (and/or un-reposited) if these hosts are natural, intermediate, or dead-end hosts. Knowledge of these factors will enable us, in future studies, to predict on these types of interactions between virus and host and hence better enable us to assess likelihood of a susceptible host being a long-term reservoir. However, the available data are too limited and too inaccessible for a study with the breadth of interactions we characterise here, and hence were unable to be included.
- Our knowledge of hosts of poxviruses remains biased towards those that infect humans and domesticated animals. To address this, we integrated research effort directly into the training phase, by assigning, to each possible virus/animal combination, a weight derived from the research effort into both the poxvirus and the animal. This enabled the underlying models to treat each virus/animal combination differently.
- Validation. Given the severe imbalance between observed associations and all potential associations between poxviruses and the animals included in this study, our top 25 ensembles performed well, detecting 93.37% of all observed associations (>0.5, 83.97 ≥0.76). Most importantly, when measured using metrics sensitive to class imbalance PR-AUC (0.485), and F1-Score (0.309 - 0.422) – our models did not inflate predicted associations, and are one of the most conservative approaches to date(52,66). However, the ideal validation would be post hoc field confirmations/observations of new interactions which we have predicted here. This is a laborious task, and beyond the scope of this work, however, the combined global research effort will continue

to identify new virus-host interactions and hence, as these data become available, our predictions may be tested.

- Note: since our data were analysed, we are aware of a single new MPXV-host association: with the domestic dog(67). Our pipeline did predict this interaction (Supplementary Dataset 1), and it represents the first of such post hoc validation possibilities.

**Supplementary Results 1 – Results for humans and domesticated animals**

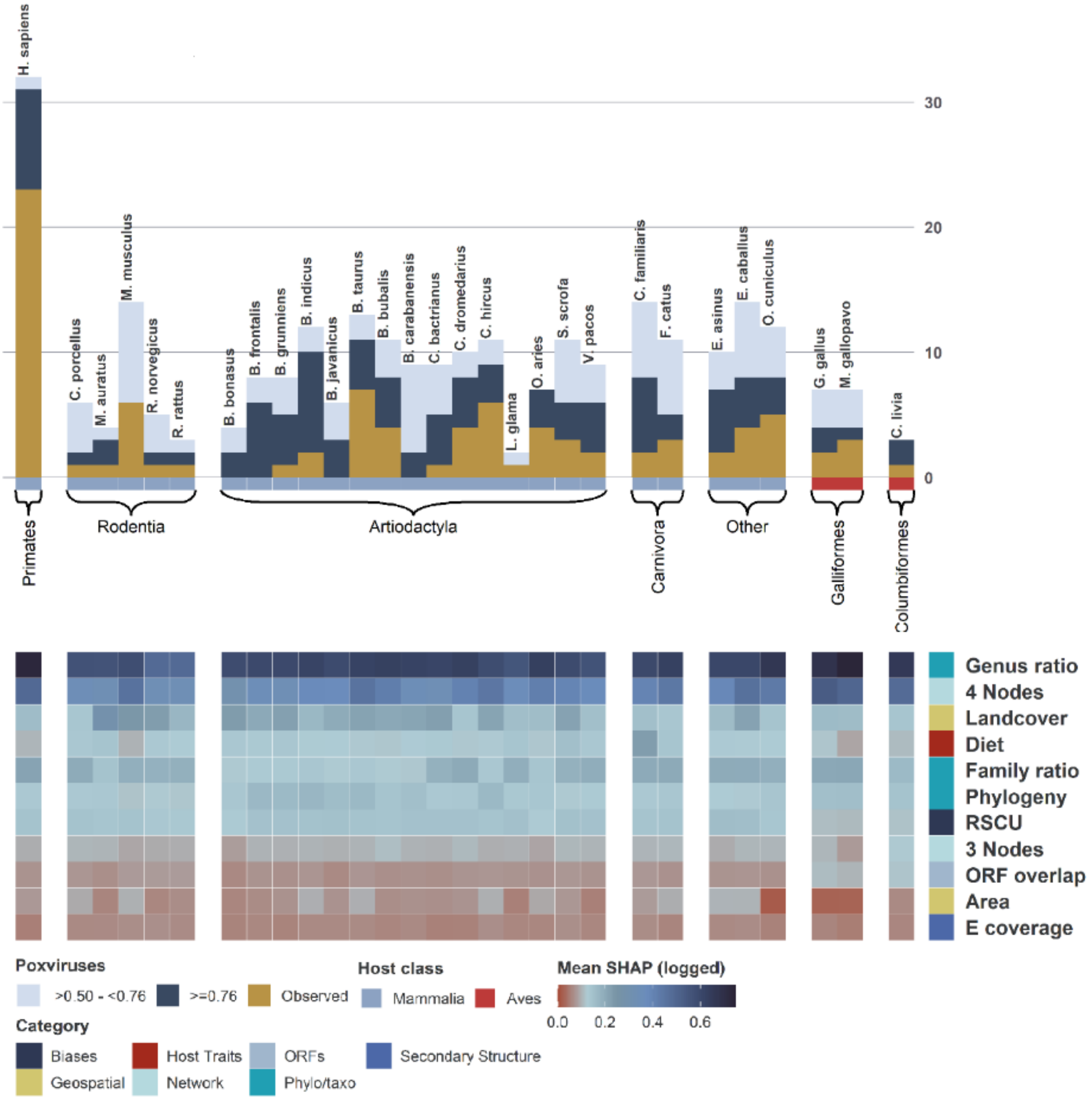


**Supplementary Figure 4 – Human and domesticated hosts.** Stacked bars represent umber of poxviruses per human and domesticated species. Species were grouped by class (mammalian or avian) and order. Yellow bars represent number of poxviruses observed to be found in each species. Blue bars show other poxviruses predicted to be found in each species by our pipeline. Predicted poxviruses per host are grouped by association probability into two categories: dark blue = ≥0.76, and light blue = >0.5 - <0.76. Heatmaps represent log10 mean absolute SHAP values. SHAP values were computed separately for each possible association of each of the included species and poxvirus (n=63), for each constituent model (n=4), of our top ensembles (n=25). Mean absolute SHAP values were taken for each association/feature combination, and then summed for each association/group combination. Features are colour coded by their category.

**Supplementary Results 2 – Order level maps**

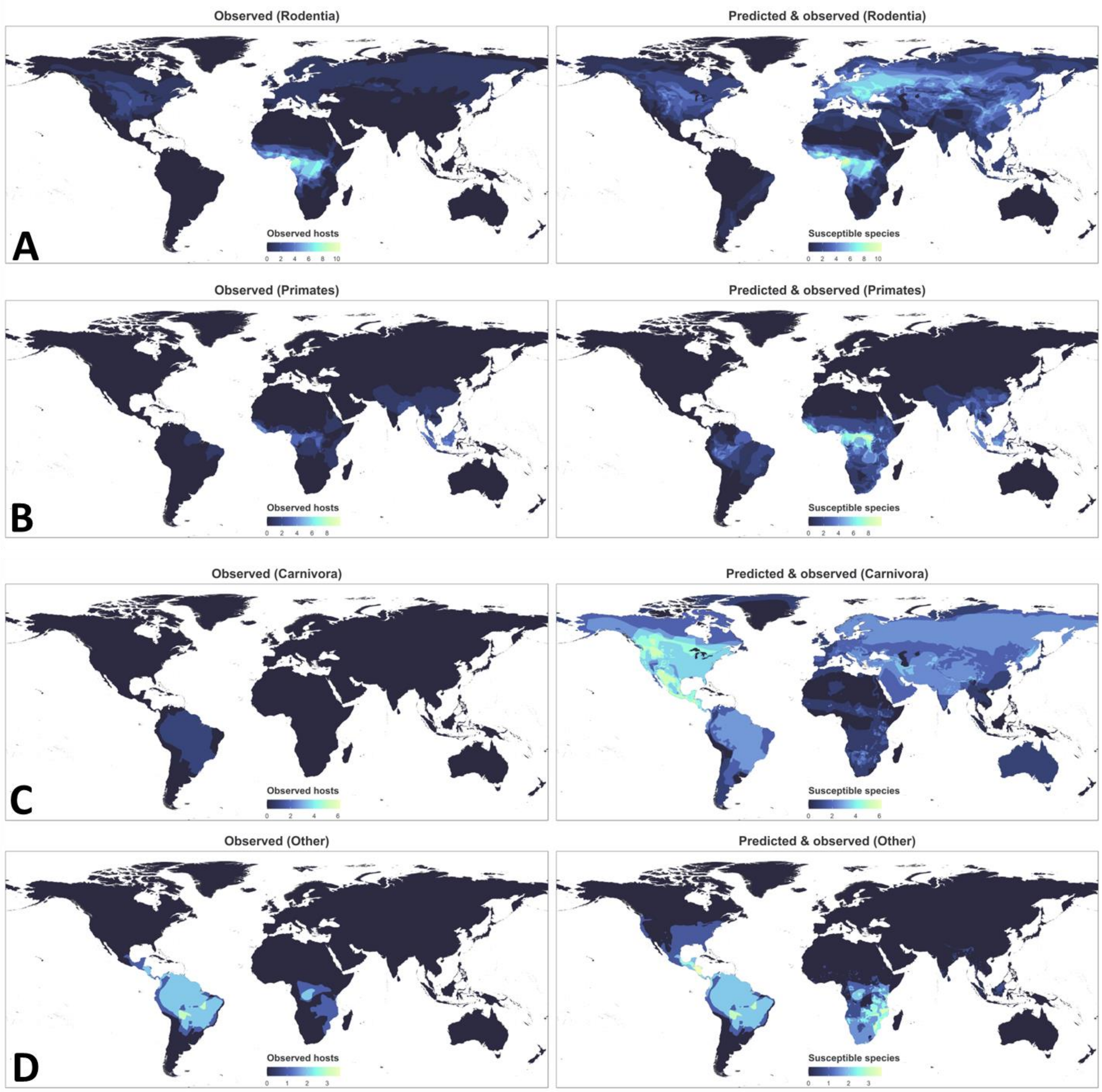


**Supplementary Figure 5 – Global distribution of observed and predicted susceptible wild terrestrial species of Monkeypox virus by order. Panel A – Rodentia. Panel B – Primates. Panel C – Carnivora. Panel D – Other mammalian orders (Artiodactyla, Didelphimorphia, Eulipotyphla, Macroscelidea, Perissodactyla, Pilosa, and Proboscidea).** Rasters were computed from presence shapefiles of observed (known) species (left-hand side) by summing number of species per each grid cell (each measuring cells measuring 1/6 x 1/6 of a degree). The right-hand side shows both observed host species and predicted susceptible species (threshold >0.5). Each grid cell is coloured by the sum of probabilities of all predicted hosts (observed host = 1; predicted host = square root of prediction value). The colour scales for both maps in each row is the same to allow comparison.

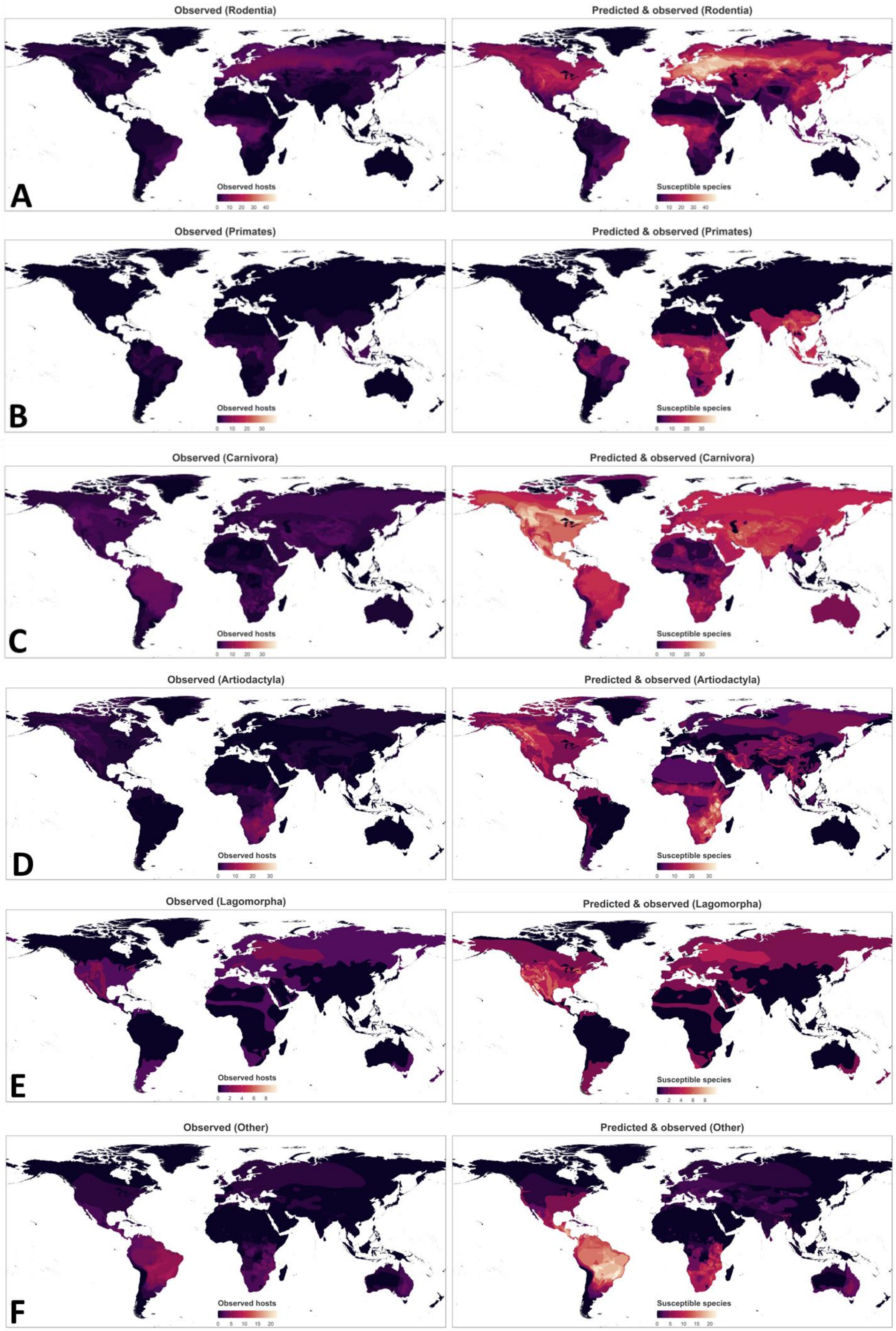


**Supplementary Figure 6 – Global distribution of observed and predicted susceptible wild terrestrial mammalian species of poxviruses by order. Panel A – Rodentia. Panel B – Primates. Panel C – Carnivora.**

**Panel D – Artiodactyla. Panel B – Lagomorpha. Panel C – Other mammalian orders (Chiroptera, Cingulata, Didelphimorphia, Diprotodontia, Eulipotyphla, Macroscelidea, Perissodactyla, Pilosa, and Proboscidea).** Rasters were computed from presence shapefiles of observed (known) species (left-hand side) by summing number of species per each grid cell (each measuring cells measuring 1/6 x 1/6 of a degree). The right-hand side shows both observed host species and predicted susceptible species (threshold >0.5). Each grid cell is coloured by the sum of probabilities of all predicted hosts (observed host = 1; predicted host = square root of prediction value). The colour scales for both maps in each row is the same to allow comparison.

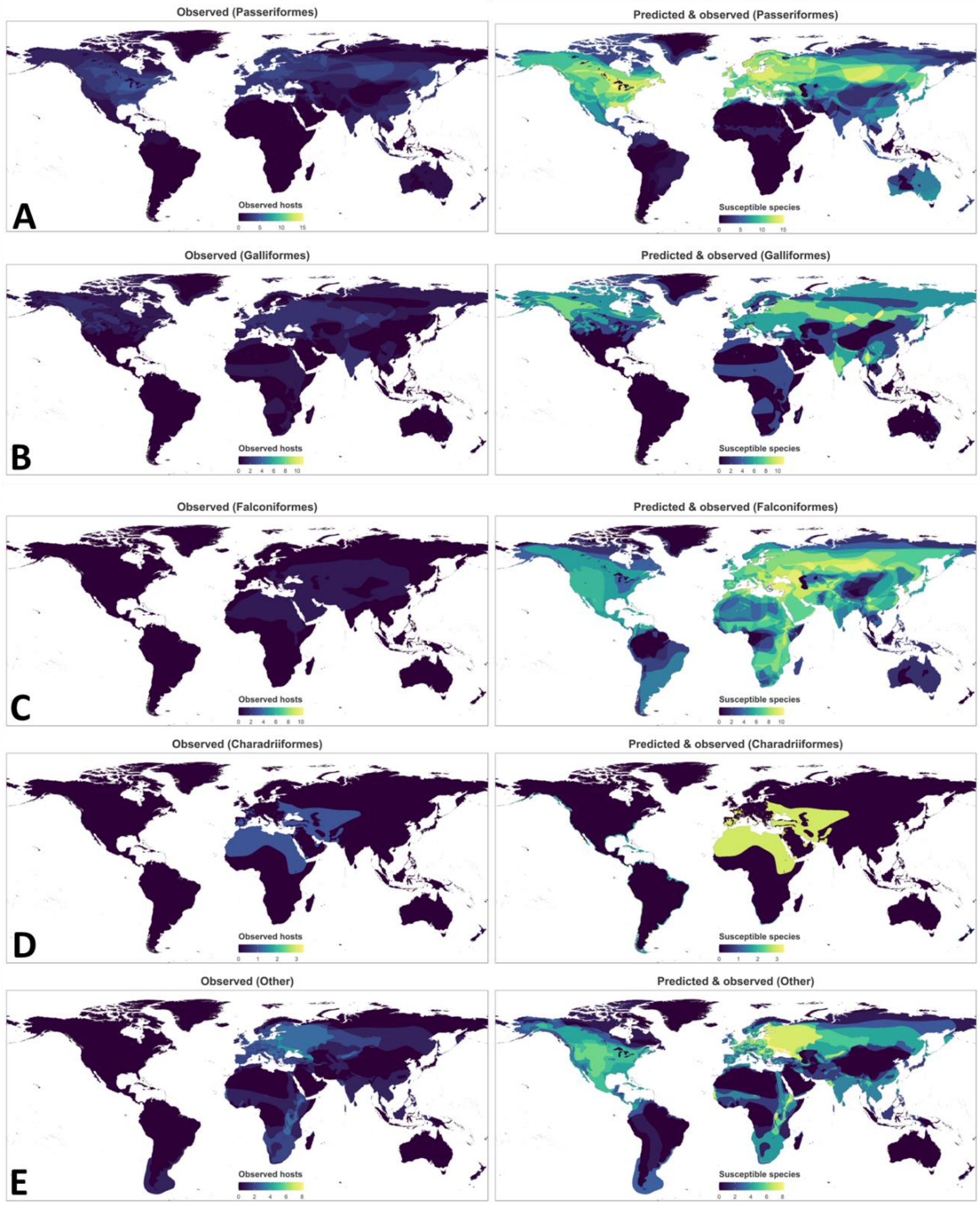


**Supplementary Figure 7 – Global distribution of observed and predicted susceptible wild terrestrial avian species of poxviruses by order. Panel A – Passeriformes. Panel B – Galliformes. Panel C – Falconiformes. Panel D – Charadriiformes. Panel E – Other avian orders (Accipitriformes, Anseriformes, Columbiformes, Gruiformes, Phoenicopteriformes, Procellariiformes, and Sphenisciformes).** Rasters were computed from presence shapefiles of observed (known) species (left-hand side) by summing number of species per each grid cell (each measuring cells measuring 1/6 x 1/6 of a degree). The right-hand side shows both observed host species and predicted susceptible species (threshold >0.5). Each grid cell is coloured by the sum of probabilities of all predicted hosts (observed host = 1; predicted host = square root of prediction value). The colour scales for both maps in each row is the same to allow comparison.

**Supplementary Results 3 – Model performance**

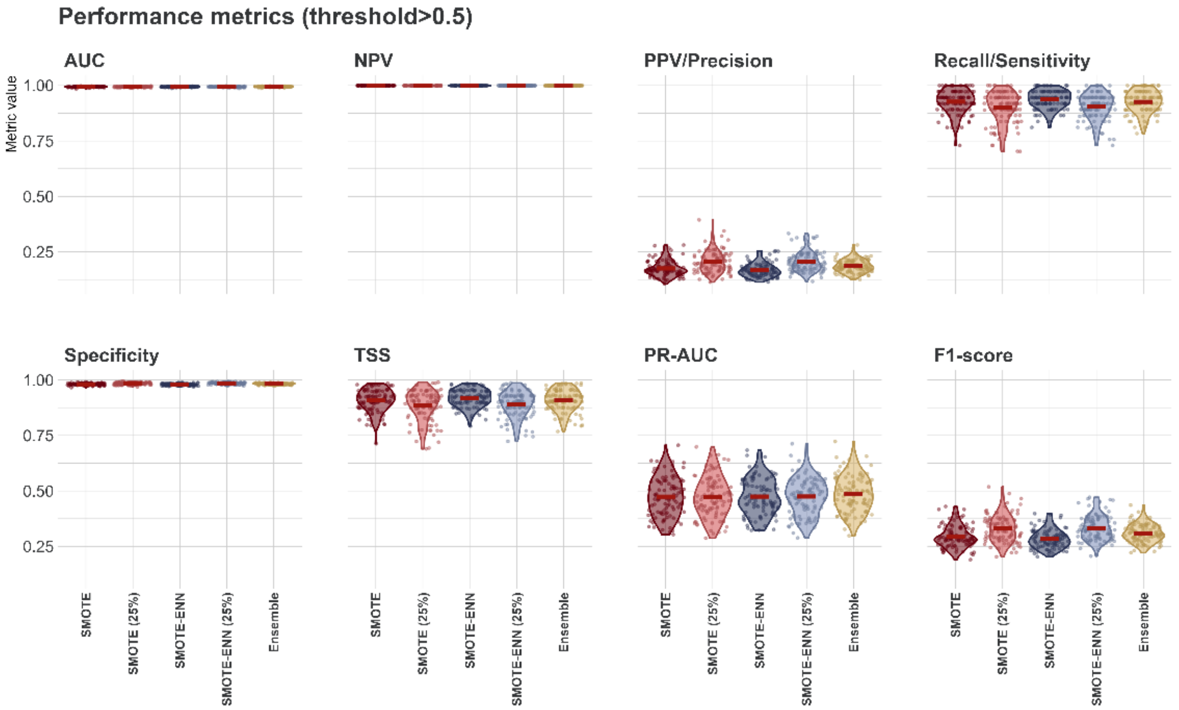


**Supplementary Figure 8 – Performance assessment over all held-out test sets (n =100) at >0.5 probability threshold.** Points represent results from individual iterations (n=100). Violin plots show the kernel probability density of the data at different values. Points and violin plots are coloured by class balancing algorithm: SMOTE, SMOTE (25%), SMOTE-ENN, and SMOTE- ENN (25%), and Ensemble (mean of the 4 algorithms). Each held-out test-set comprise a 10% stratified random sample of all possible associations (n= 87,532).

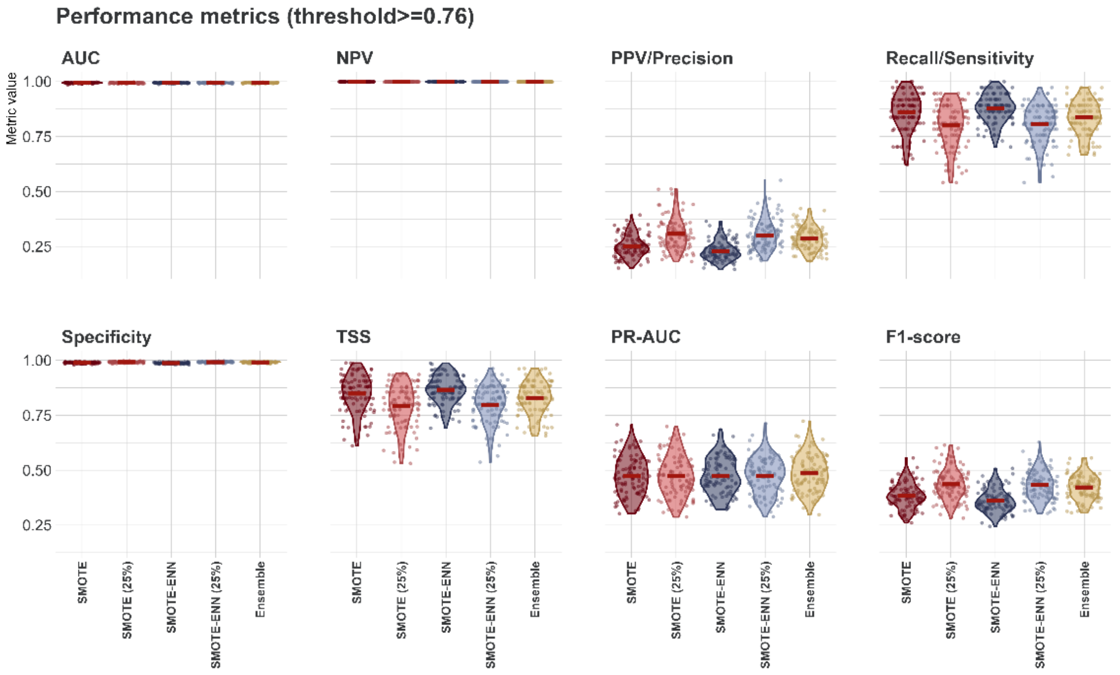

**Supplementary Figure 9 – Performance assessment over all held-out test sets (n =100) at ≥0.76 probability threshold.** Points represent results from individual iterations (n=100). Violin plots show the kernel probability density of the data at different values. Points and violin plots are coloured by class balancing algorithm: SMOTE, SMOTE (25%), SMOTE-ENN, and SMOTE- ENN (25%), and Ensemble (mean of the 4 algorithms). Each held-out test-set comprise a 10% stratified random sample of all possible associations (n= 87,532).

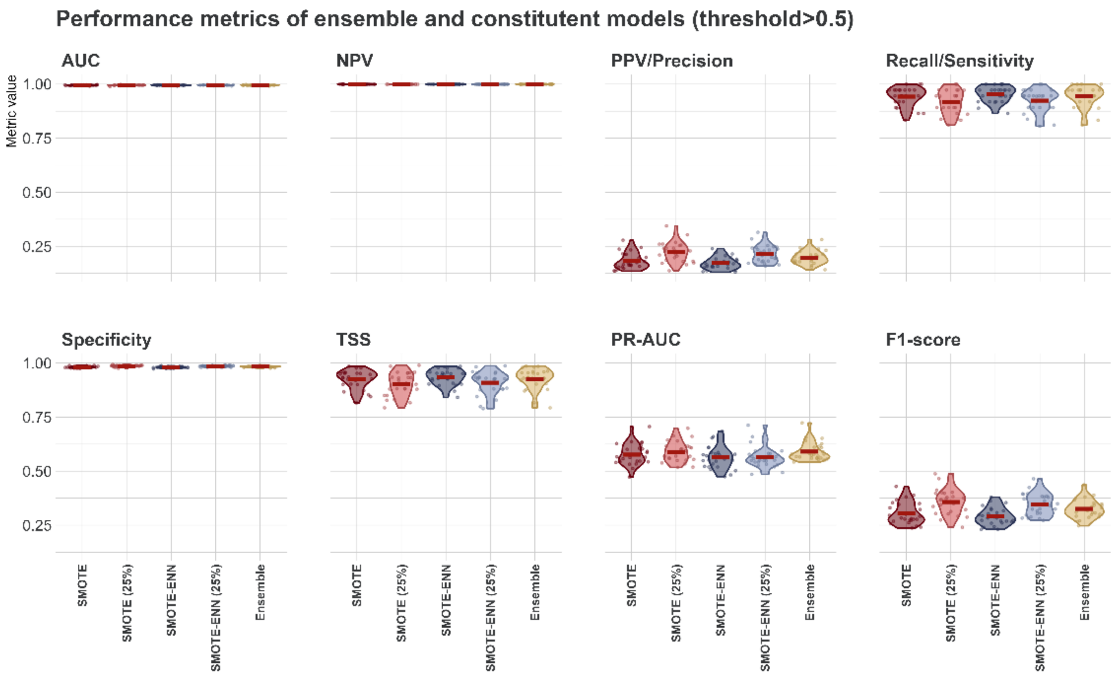


**Supplementary Figure 10 – Performance assessment over held-out test sets of selected top 25 ensembles at >0.5 probability threshold.** Points represent results from individual iterations (n=25). Violin plots show the kernel probability density of the data at different values. Points and violin plots are coloured by class balancing algorithm: SMOTE, SMOTE (25%), SMOTE-ENN, and SMOTE- ENN (25%), and Ensemble (mean of the 4 algorithms). Each held-out test-set comprise a 10% stratified random sample of all possible associations (n= 87,532).

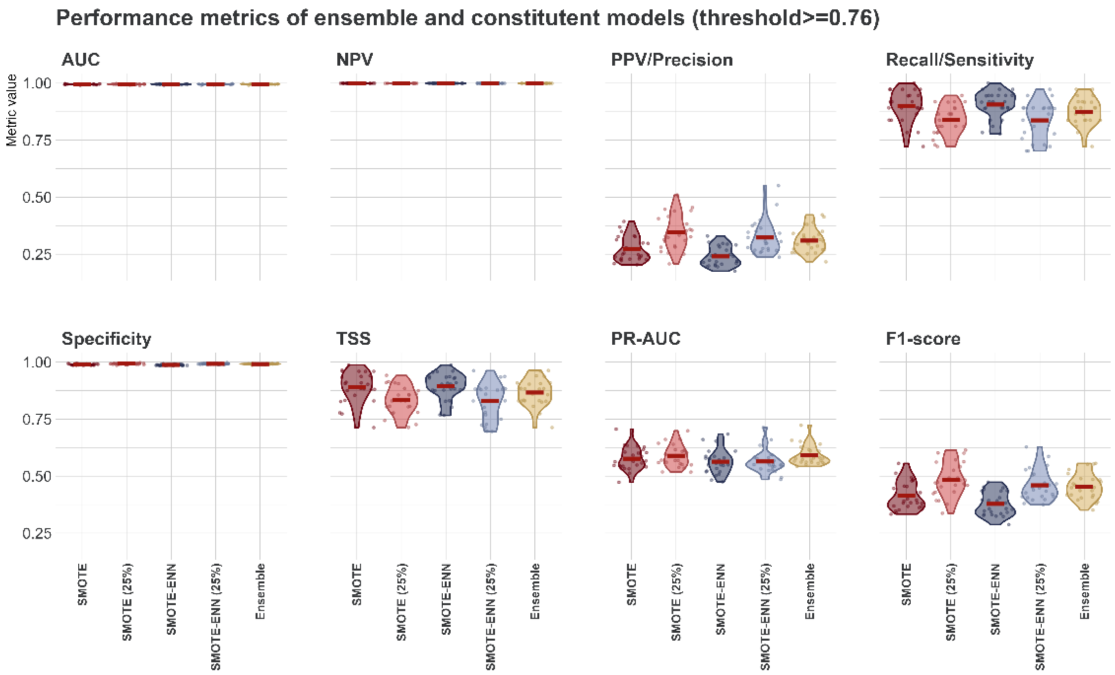


**Supplementary Figure 11 – Performance assessment over held-out test sets of selected top 25 ensembles at ≥0.76 probability threshold.** Points represent results from individual iterations (n=25). Violin plots show the kernel probability density of the data at different values. Points and violin plots are coloured by class balancing algorithm: SMOTE, SMOTE (25%), SMOTE-ENN, and SMOTE- ENN (25%), and Ensemble (mean of the 4 algorithms). Each held-out test-set comprise a 10% stratified random sample of all possible associations (n= 87,532).
